# Emotional Metacognition: Stimulus Valence Modulates Cardiac Arousal and Metamemory

**DOI:** 10.1101/2020.06.10.144428

**Authors:** Nicolas Legrand, Sebastian Scott Engen, Camile Maria Costa Correa, Nanna Kildahl Mathiasen, Niia Nikolova, Francesca Fardo, Micah Allen

## Abstract

Emotion alters how we feel, see, and experience the world. In the domain of memory, the emotional valence and arousal of memorised stimuli can modulate both the acuity and content of episodic recall. However, no experiment has investigated whether arousal and valence also influence metacognition for memory (i.e., the process of self-monitoring memories). In a pre-registered study, we applied a novel psychophysiological design together with computational models of metacognition to assess the influence of stimulus valence and arousal on the sensitivity, bias, and efficiency of metamemory. To estimate the role of physiological arousal in mediating these effects, we recorded cardiac measures through pulse oximetry. We found that negative valence substantially decreased both memory performance and subjective confidence, in particular for low arousal words. Simultaneously, we found that emotional valence modulated both heart rate and heart-rate variability (HRV) during recognition memory. Exploratory trial-level analyses further revealed that subjective confidence was encoded in instantaneous heart-rate fluctuations and that this relationship was also modulated by emotional valence. Our results demonstrate that recognition memory and metacognition are influenced by the emotional valence of encoded items and that this correlation is in part related to cardiac activity.

## Introduction

The cognitive ability to monitor our thoughts, memories and perceptual experiences is an important part of learning, development and communication (Fleming et al., 2012; Heyes et al., 2020; Shea et al., 2014). Metacognition refers to this executive capacity to control lower-order representations by assessing the fidelity and strength of these signals and updating a model of the probability that one is making correct judgements (Yeung & Summerfield, 2012). More specifically, metacognition for memory or metamemory refers to the ability to assess the accuracy and precision of the recollection of previous experiences (Baird et al., 2013; McCurdy et al., 2013; Ye et al., 2018). Memories are fragile internals signals that are prone to substantive decay over time (Davis & Zhong, 2017; Otgaar et al., 2019). Episodic recall can be biased by emotion and arousal both in the context of encoding (Yonelinas & Ritchey, 2015) and during active recall (Ochsner, 2000). The metacognition of memory recall can, in turn, be influenced by the level of details and the “feeling-of-knowing” associated with an episode (Chua et al., 2014; Reggev et al., 2011). In the context of witness testimony for example, if the suspect had an unremarkable face or the memory was hazy, then a witness may report lower confidence in their recollection. A reliable witness should therefore not only recall events as experienced in detail but also accurately assess the fidelity or confidence associated with those memories. As little is currently known about the ability to self-monitor memory for emotional stimuli, we conducted a confirmatory, pre-registered investigation of emotional metamemory.

In controlled laboratory settings, classic metacognition experiments often require participants to view a stimulus, make a decision (e.g., whether the stimulus is known or unknown), and report their confidence in this judgement. Healthy individuals typically display reasonably accurate metacognitive insight and achieve a high correlation between confidence and accuracy, even in the absence of external feedback. Metacognition tasks have been applied to investigate a variety of cognitive domains including visual perception (Allen et al., 2016; Fleming et al., 2015), memory (Baird et al., 2013; Fleming et al., 2014; McCurdy et al., 2013), and value-based decision-making (De Martino et al., 2013). However, even with these simple lab-based tasks, participants exhibit substantive interindividual differences in metacognitive ability, and a variety of manipulations can reliably dissociate confidence and accuracy by biasing subjective confidence reports (Fleming et al., 2015; Rollwage et al., 2020).

Though little is known about how emotion influences metacognition, previous investigations of memory and emotion highlight stimulus valence and arousal as likely sources of bias. For example, words rated as highly arousing or positively valenced are recognized and detected faster (Kever et al., 2019), and the emotional content of valenced words, either positive or negative, can increase the learner’s confidence in subsequent accurate recall (Tauber & Dunlosky, 2012). Similarly, physiological arousal can promote the awareness of concepts congruent with the bodily state (Kever et al., 2015, 2017), and the level of arousal at encoding can enhance subsequent memory performance (Anderson, 2005; Cahill & McGaugh, 1998) via greater amygdala activity (Kensinger & Corkin, 2004). Accordingly, flashbulb memories (i.e., vivid and detailed memories encoded under arousing conditions) are recalled more easily and with less decay under specific circumstances (Shields et al., 2017; Yonelinas & Ritchey, 2015). This line of evidence suggests that emotional content, especially those of a highly arousing or negative nature, could bias the salience of the memory signal during recall. This can ultimately result in overconfidence which, in the context of testimony, could bias the individual when estimating the accuracy of his or her recall.

Additionally, a core aspect of emotion is that it often coincides with and is triggered by changes in internal bodily states (James, 1884), which is expressed by indices of autonomic activity such as cardiac or respiratory frequency (Kreibig, 2010; Valenza et al., 2014). Heart rate, for example, can be altered both when perceiving emotional stimuli and during their encoding and recollection (Abercrombie et al., 2008; Critchley et al., 2005; Legrand et al., 2020). These bodily changes exert a substantial effect on the mapping between confidence and decision accuracy, which can ultimately also bias metacognition. For example, both experimental and pharmacological modulations of arousal have been shown to bias metacognitive insight, modulating confidence for error trails in a visual task (Allen et al., 2016; Hauser et al., 2017).

Here, we hypothesized that both the valence and arousal of an encoded stimulus might modulate the accuracy of memory itself, and investigated whether healthy individuals are aware of such emotional effects on their recognition accuracy. To test this hypothesis, we conducted a pre-registered experiment in which participants memorised lists of words grouped by their levels of valence and arousal. Although most metamemory research has relied on “feeling of knowing” self-report measures, these can be subject to substantive biases, i.e. such as conflating self-report bias with metacognitive sensitivity or being confounded by overall accuracy level (Fleming & Lau, 2014). To overcome these issues, we adapted a signal-theoretic modelling approach to estimate recognition metamemory for emotional versus unemotional words. If arousal primarily biases memory, for example by increasing the salience of encoded items, we would expect to observe a positive main effect of item arousal on both accuracy and metacognitive confidence. Conversely, if emotion primarily biased metacognition through a valence-specific ‘anchoring’ effect, we would expect to observe a full interaction of stimulus arousal and valence on both measures. As a third alternative, if metacognition were robust to emotional biases, we would expect to observe the effects of stimulus valence and arousal on accuracy and response speed, but not on confidence or metacognition. To complement these analyses, we further recorded cardiac measures of physiological arousal through pulse oximetry, to assess their mediating effect on the association between confidence and accuracy.

## Materials and methods

### Pre-registration and Open Materials

To improve our control of type-I and type-II error rates, as well as the overall reproducibility of the study, the trial was pre-registered before any data collection using the standard Open Science Foundation template. Detailed information regarding power analysis, sample size considerations, experimental and trial design, planned analyses and other key points can be found at the following URL: (https://osf.io/9awtb). In what follows, Confirmatory Analyses and Results refer to planned analyses detailed in the pre-registration, whereas Exploratory Analyses and Results refer to post-hoc exploratory analyses conducted following contact with the data. Additionally, in the case of any minor deviation from the pre-registration, these are documented on a case by case basis.

### Participants

Thirty-five participants (26 females) between the ages of 18 and 26 (M = 21, SD = 1.9) were recruited through local advertisements and took part in the experiment at Aarhus University Hospital, Denmark. From the total sample of 35 participants, a sub-set of 30 participants passed the pre-registered exclusion criteria and were analyzed further. All participants had normal or corrected to normal vision, were fluent in English and provided informed written consent before the experiment. The procedures were conducted following the Declaration of Helsinki and with approval from the Danish Neuroscience Centre’s (DNC) Institutional Review Board (IRB). Participants received monetary compensation of 100 DKK per hour. The estimated total duration of the test session was 1,5 hours (150 DKK). Participants also completed a post-test stimulus validation measure in which they provided valence and arousal ratings for all stimuli for an additional 50 DKK. All 35 participants completed the follow-up rating experiment.

### Procedure

The experimental procedure included one laboratory and one at-home survey session on two different days with one week in between. In the laboratory session, participants completed a word recognition metamemory task designed to assess the effects of valence and arousal on verbal recognition memory and metacognition. In the survey session, participants rated their subjective feelings of valence and arousal evoked by the words used during the laboratory session.

At the beginning of the laboratory session, participants were briefed on the nature of the investigation, were provided task instructions and completed a brief training session of the metamemory task. The training included an example learning phase of 50 neutral and unarousing words, followed by an example testing phase of 10 trials with confidence ratings (see Metamemory Task and Stimuli).

During the metamemory task, heart rate was monitored using a Nonin 3012LP Xpod USB pulse oximeter together with a Nonin 8000SM ‘soft-clip’ fingertip sensor (https://www.nonin.com/) attached to the left index finger.

### Word selection

Stimuli consisted of 1200 English words selected from the Affective Norms for English Words (ANEW) database based on valence and arousal ratings measured among a population of American students (Bradley & Lang, 1999). Although ANEW is not validated in the Danish population, previous standardization of the database in Dutch, Spanish and Italian populations (Montefinese et al., 2014; Moors et al., 2013; Redondo et al., 2007) showed good consistency across both American and European samples. In order to estimate the reliability of these ratings among our sample, we conducted an at-home valence and arousal rating task after the behavioural test. We created 4 distinct subgroups of 300 word stimuli, according to a 2 by 2 factorial design, where the factors corresponded to valence (positive vs. negative) and arousal (low vs. high). To this aim, we used the tertile of the valence and arousal distribution to exclude words with intermediate ratings, whose valence and arousal might be ambiguous (see Fig. 2a).

### Metamemory Task

Participants completed a word recognition metamemory task adapted from a previous study (McCurdy et al., 2013) to test the influence of emotional valence and arousal on memory and metacognition. The task included 12 blocks, each consisting of a learning phase (Fig. 1a) and a testing phase (Fig. 1b). In the learning phase, participants viewed a list of 50 English words for durations of 30, 60 or 90 seconds. The words were presented on the screen in the form of a table containing five columns with ten rows of words each, and the participants were instructed to memorize as many words as possible. The list of words in each learning phase corresponded to a unique combination of the following factorial conditions: Valence (positive, negative), Arousal (low, high). Participants were notified when 10 seconds of learning time was left by the display of a small warning at the bottom of the screen. During the testing phase, participants completed 50 trials designed to measure recognition memory and metamemory. On each trial, two-word stimuli were presented to the left and right of a fixation cross. The word pair consisted of a “target” and a “distractor”, corresponding to words that were present or absent in the previous learning phase, respectively. Target and distractor words were matched by valence and arousal, and their position was randomized across trials. Participants were instructed to press either the left or the right arrow key to indicate which of the two words they recognized from the memorised list. This procedure corresponds to a two-alternative forced-choice task (2AFC) design, which provides optimal conditions for estimating and comparing metacognition scores across tasks (Lee et al., 2018). Following the button press, participants provided a subjective confidence rating from 1 (“not confident at all/guessing”) to 7 (“very confident”). Both button presses and confidence ratings had a maximum time-limit of 3s. If participants had slower responses, a brief message (i.e., “too slow!”) was displayed on the screen and the trial was marked as missed.

**Figure 1:**
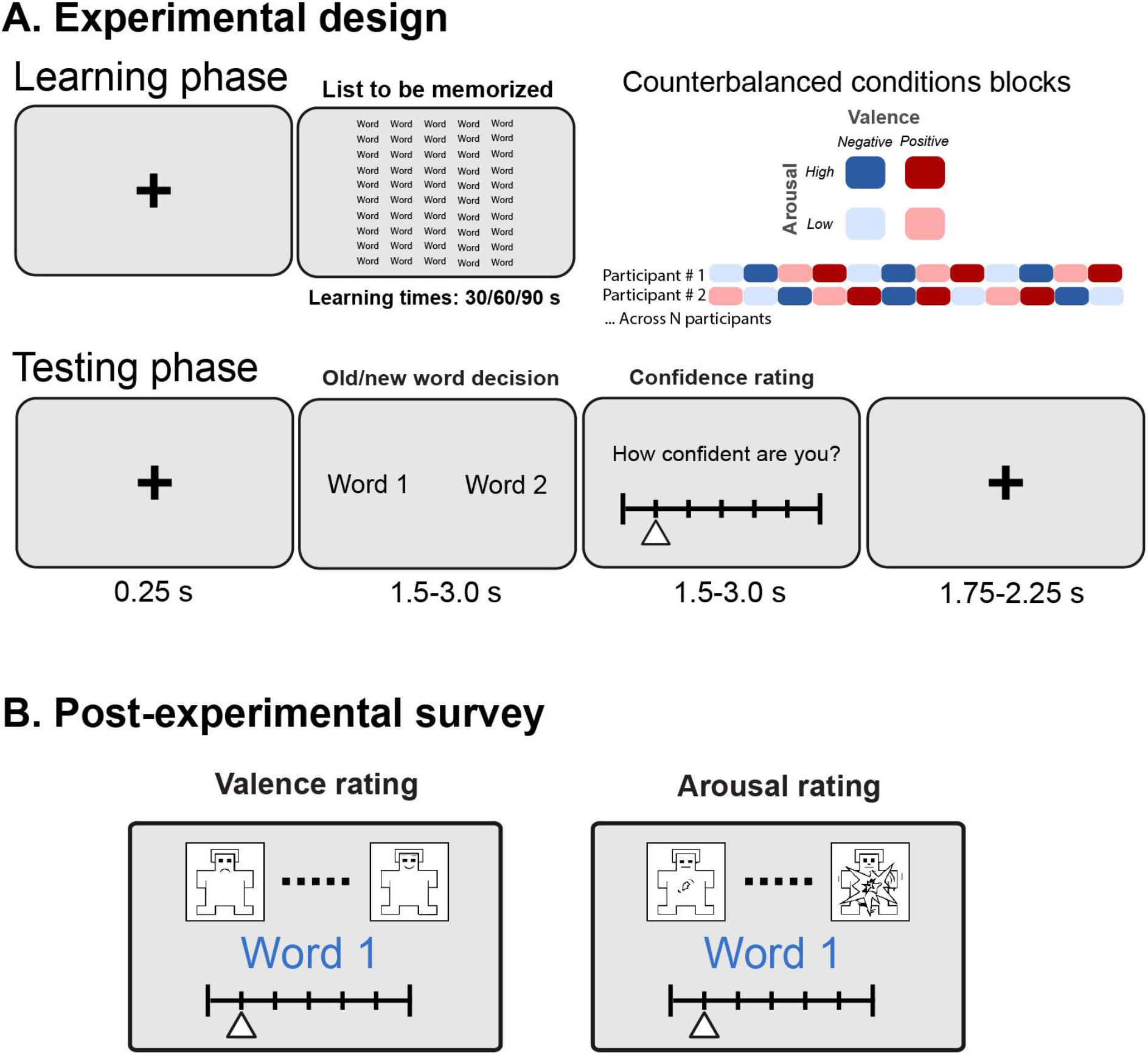
**A.** Experimental design. The metamemory task contained 12 experimental blocks, each consisting of a learning phase and a testing phase for the 50 words, in a factorial design separated by each Valence and Arousal condition. To limit habituation effects, block orders were counter-balanced in a pseudo-randomized order such that each high arousal block was interceded by a low arousal condition. **B.**Post-experimental survey. To validate our stimulus categories with respect to the original ANEW ratings, participants completed a short subject visual analogue scale rating of valence and arousal for the 1200 words used in the main task (600 target and 600 distractors) in an at-home experiment. This was done in a web-based version of the original procedure used in the original ANEW survey.

The blocks were presented in a pseudo-randomized order to ensure that high arousal blocks were systematically interleaved with low-arousal blocks. The block order and the selection of target vs. distraction lists were counterbalanced across participants.

### Valence and Arousal Rating Task

To validate the arousal and valence stimulus categories in our Danish sample, participants completed an at-home valence and arousal subjective rating task. They were instructed to provide valence and arousal ratings of their subjective experience associated with each word presented in the metamemory task. The ratings were collected using a 9-point visual numerical scale in a web-based version of the original ANEW survey protocol (Bradley & Lang, 1999). Our version was implemented using Pavlovia (https://pavlovia.org), an online platform for running PsychoPy experiments (Peirce et al., 2019). Each word was presented twice, once for valence and once for arousal, and the 9-point scales were complemented with pictures of the original drawings of the Self-Assessment Manikin (Bradley & Lang, 1994), as in the original ANEW survey (Bradley & Lang, 1999). Participants rated a total of 1200 words, self-pacing through all rating trials. We compared the ratings provided by the participants in this study with the normative ANEW ratings using a Spearman rank correlation test (see Fig. 2 c & d). After inspecting histograms of participant responses, we excluded one participant, who only ever pressed the same key (rating 5); this exclusion criterion was not noted in the pre-registered protocol. Overall, stimulus ratings in our sample corresponded very well to the original ANEW ratings, ρ = [0.93-0.78], albeit with lower overall consistency for the arousal vs. valence dimension (Fig. 2).

**Figure 2:**
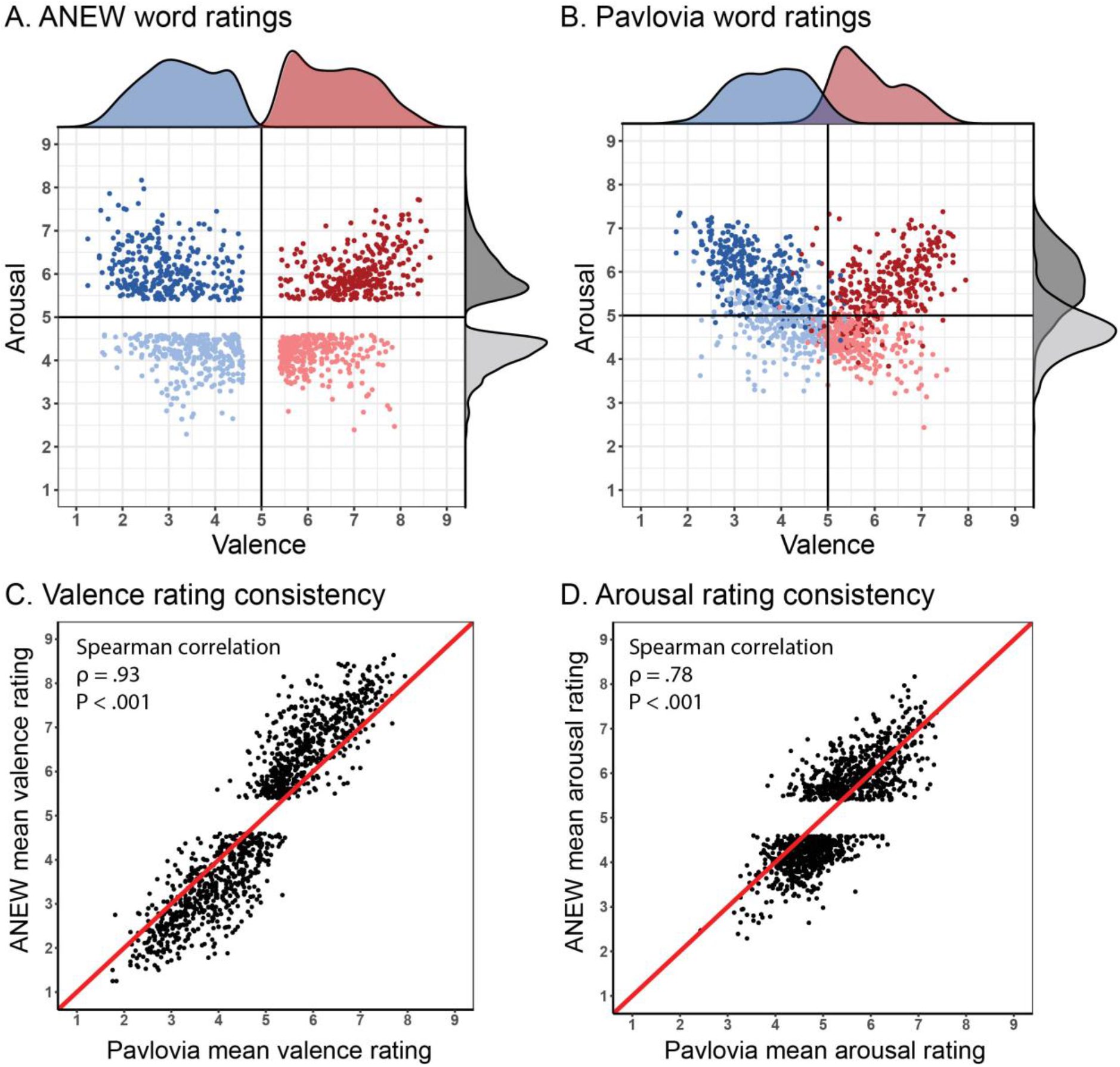
Stimulus selection and rating validation. Two rating procedures were used to select and validate word stimuli; the original ANEW normed ratings and the PAVLOVIA at-home ratings completed by our participants in the post-experimental survey. Each word was rated on a 9-point scale (1-9) for valence and arousal separately. **A & B**. We selected the words used in the metamemory task by removing items from the central tertile in the arousal and valence rating distributions (panel A). The blue and red dots represent words with negative and positive valence, respectively. The light and dark points represent low and high arousal, respectively. The densities represent the distribution for positive and negative valence (red and blue), and arousal (light and dark). **C & D.**We compared the independent rating provided by the ANEW database to the actual ratings provided by the participants, PAVLOVIA, after the main procedure. Both ratings of valence and arousal showed reasonably high consistency, ρ = [0.93-0.78]. See online article for colour figures. The black dots represent each word in the datasets and the red line shows the identity line.

**Fig. 3.**
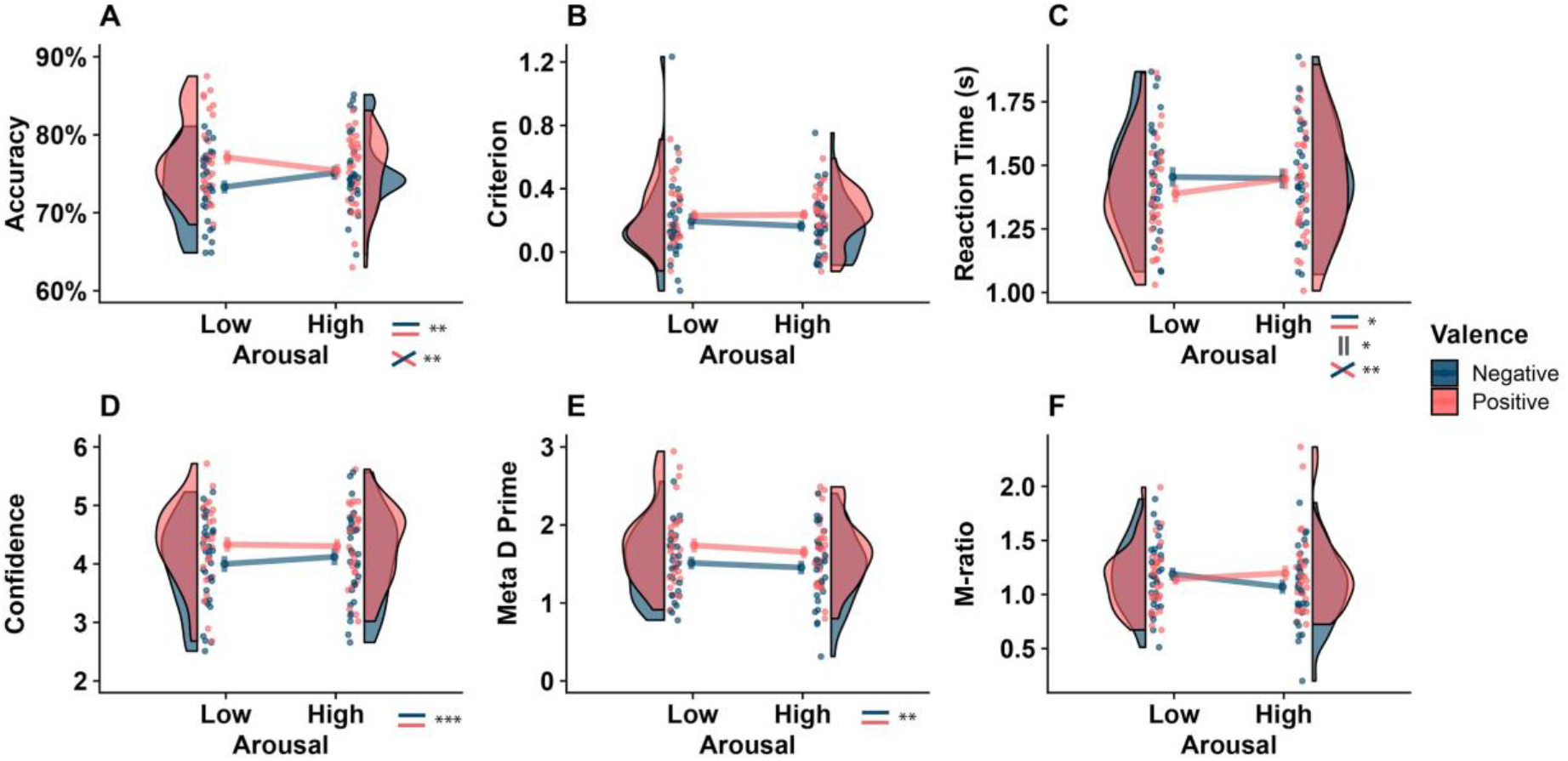
Behavioural results showing factorial main effects and interactions on discrimination and metacognitive performance. Modified raincloud plots (Allen et al., 2019) illustrating behavioural results of discrimination measures of accuracy (**A**), criterion (**B)** and reaction time (**C**) as well as metacognitive measures of confidence (**D**), Meta-d’ (**E**) and M-ratio (**F**). Repeated measures ANOVA (Valence × Arousal) was carried out for each condition separately. The upper panel shows that a significant main effect of emotional valence was observed as negative valenced words reduced accuracy (**A**) and slowed down reaction times (**C**). Similarly, the lower panel shows a main effect of valence for both Confidence and Meta-d’ are impaired by negative valence. (*** p<0.001, ** p<0.01, * p<0.05).

### Signal Theoretic Metacognition Modelling

Here we applied a signal-theoretic computational model to describe participant behaviour on the metamemory task (Fleming, 2017; Maniscalco & Lau, 2012). This approach delineates overall behaviour into ‘type-I’ and ‘type-II’ measures, corresponding to a basic discrimination performance versus metacognitive levels of performance (Galvin et al., 2003). The type-I performance was quantified using reaction times (RTs) and the signal-detection theoretic (SDT) measures of d-prime (D’) and criterion (c) (Macmillan & Creelman, 2004). Type-II performance (i.e., metacognition) was assessed by the SDT measures of Meta-d’ and Meta-ratio (M-ratio) (Fleming & Lau, 2014). Briefly, this approach models metacognitive “hits” (e.g., high/low confidence for correct/error trials, respectively) and “misses” (e.g., high/low confidence for error/correct trials, respectively). All SDT-based measures were estimated at the subject level (Fleming, 2017). This model has been extensively described and validated previously (Fleming, 2017; Mazancieux et al., 2020; Morales et al., 2018), here we recount the approach in the context of the present study to aid interpretation.

D-prime or d’ is a measure of a participant’s sensitivity to detect previously studied words during the learning phase, independently from subjective response biases. Instead, criterion or c’ encodes the participant’s response bias, that is, the overall tendency to prefer one response over the other (e.g. if a participant chose the word presented to the left of the fixation point more often than the alternative). Together with measures of reaction time, d’ and c’ are metrics of “first-order” or “type-I” task performance. In contrast, Meta-d is an estimate of the sensitivity of subjective confidence ratings to type-I performance (i.e., the probability to be highly confident when correct, or uncertain when incorrect). Meta-d’ is, therefore, a measure of insight, or how well one can consciously discriminate their own type-I performance (Lau & Rosenthal, 2011). However, metacognitive sensitivity is also a function of the overall perceptual signal, and as such is substantively influenced by differences in d’. To control for this effect, the ‘M-ratio’ (Meta-d’ divided by d’) is estimated as a measure of metacognitive efficiency, denoting how a subject’s metacognitive sensitivity over- or underperforms what can be expected given their type-I sensitivity (Fleming & Lau, 2014). Finally, average confidence on each condition denotes participants “meta-criterion” or “meta-c”, or their overall level of metacognitive bias denoting the tendency to be confident or uncertain irrespective of accuracy. Meta d’ and meta-c are metrics of “second-order” or “type-II” performance.

### Confirmatory Analyses

#### Metamemory task

All data were pre-processed according to the protocols established in our pre-registration. Accordingly, we excluded all trials with reaction times (RT) faster than 100 ms, greater than 3 standard deviations from the median RT, and missing data (absence of response or because the response button was pressed too early or too late). Due to an unforeseen technical error, an absence of response in some trials contaminated the following trial, resulting in negative response times. These trials were also automatically rejected. This procedure resulted in the exclusion of 3.49% (±3.68) of the trials. Finally, outliers in task performance for each of the conditions were detected based on reaction time, d’, and confidence distributions. We also excluded participants showing any extreme value using Tukey’s boxplots. Based on these criteria, 5 participants were excluded from all behavioural analyses. These preprocessing steps are also extensively described in the interactive Jupyter notebooks made available on the OSF repository: https://osf.io/9awtb.

The preprocessing of the behavioural data was carried out using custom R scripts, using R Studio (1.2.5019), the R software (R 3.6.1), and Python scripts using Python 3.7.6. The Bayesian and frequentist statistical models were implemented using the JASP software (https://jasp-stats.org/) version 0.12.2 and the R package (AFEX 0.27-2). All Type-1 and Type-2 SDT measures (d’, criterion, meta-d’, m-ratio, and mc) were derived from the hierarchical meta-cognition model (HMM) (Fleming, 2017) implemented in R (https://github.com/metacoglab/HMeta-d), run on the individual level to enable frequentist analysis of the resultant parameters.

#### Heart Rate Monitoring

We monitored instantaneous heart rate variability using a Nonin 3012LP Xpod USB pulse oximeter together with a Nonin 8000SM ‘soft-clip’ fingertip sensor (https://www.nonin.com/). Pulse oximeters indirectly measure peripheral blood oxygen saturation. The abrupt cyclic increase of oxygenation reflects blood pulse following cardiac contraction. Here, we used the pulse-to-pulse intervals to estimate the instantaneous heart rate. Oxygenation saturation level was continuously recorded at a 75 Hz sampling rate. The preprocessing of the pulse oximetry recording was carried out using Python scripts (Python version 3.7.6) and version 0.1.1 of the Systole Python package (Legrand & Allen, 2020). Statistical analyses were carried out using the Pingouin Python package (Vallat, 2018) and MNE Python (Gramfort, 2013). PPG signals were first upsampled to 1000 Hz and clipping artefacts were corrected using spline interpolation following recent recommendations (van Gent et al., 2019). The signal was then squared for peak enhancement and normalized using the mean + standard deviation using a rolling window (window size: 0.75 seconds). All positive peaks were labelled as systolic (minimum distance: 0.2 seconds). We then detected ectopic, long, short, missed and extra beats using adaptive thresholds over the successive beats-to-beats interval (Lipponen & Tarvainen, 2019), as implemented in Systole (Legrand & Allen, 2020). The code implementing these steps can be found in the Jupyter notebooks made available on the OSF repository: https://osf.io/9awtb.

##### Instantaneous Pulse Rate

All pulses labelled as missed or extra beats were corrected by adding or removing beats, respectively. We then interpolated the instantaneous heart rate at 75 Hz to a continuous recording using the previous values and divided it into epochs (from-1 second pre-trial to 6 seconds after the word presentation). All the epochs that contained, or were adjacent to, an interval that was labelled as long, short or (pseudo-)ectopic beats were automatically rejected, resulting in an average rejection rate of 18.22% (±11.49%). The instantaneous heart rate was then averaged across trials for each condition and downsampled to 5 Hz for subsequent analyses.

##### Linear regression

In an exploratory analysis, we used the instantaneous pulse rate as a predictor of confidence over time to track the relationship between cardiac frequency modulation and metamemory. We extracted the data following the same procedure, this time using 1s before the trial start as a baseline and using the initial sampling rate (75 Hz) to facilitate cluster-based statistical tests. Cluster-based permutation testing was performed using the *permutation_cluster_test()* and the *permutation_cluster_1samp_test()* functions from the MNE Python package (Gramfort, 2013). This enabled us to assess significant point-to-point deviations from zero in encoded responses while controlling for multiple comparisons.

##### Pulse Rate Variability

Besides the analysis of the instantaneous pulse rate, we also performed pulse rate variability analyses. Although targeting a different physiological signal as compared to a classic electrocardiogram (ECG), the varying length of pulse cycle provides a sufficiently accurate estimation of the underlying heart rate variability (HRV) when used at rest for healthy young participants (Schäfer & Vagedes, 2013). Here, we extracted the systolic peak intervals using the method presented above. Intervals labelled as missed or extra beats were corrected by adding or removing beats, respectively. Additionally, intervals that were labelled as short, long, or (pseudo-)ectopic beats were corrected using linear interpolation. Following our specification in the pre-registration, we reported heart rate variability metrics in the time (RMSSD, pnn50) and frequency domain (normalized and non-normalized high and low-frequency power), as well as non-linear indexes (SD1 and SD2). These indexes reflect changes in beat-to-beat intervals and measure sympathetic and parasympathetic influences on the heart (Shaffer et al., 2014; Shaffer & Ginsberg, 2017). We inspected the resulting time series and rejected noisy and unreliable segments (2 segments were rejected in total). The values of each metric were then averaged across learning time (30, 60 or 90 seconds), and the summary variables were entered into a 2 by 2 repeated-measures ANOVA with factors stimulus arousal (arousing vs. unarousing) and valence (positive vs. negative).

## Results

### Behavioural Results

#### Overview

Following our pre-registration, the behavioural analyses focused on two levels of performance during the metamemory task: type-I variables corresponding to the discrimination ability, and type-II variables describing metacognition. To assess memory performance, we analyzed decision accuracy, discrimination sensitivity (d’), bias (c), and response time (RT). To assess metacognition, we analyzed average confidence (i.e., metacognitive bias), metacognitive sensitivity (Meta-d’), and metacognitive efficiency (M-ratio, Meta-d’/d’). All signal theoretic measures (d’, c, Meta-d’, and M-ratio) were estimated using a unified Bayesian approach, as described previously (Fleming, 2017). All posthoc tests were corrected for multiple comparisons using the Holm procedure. Here, we reported only the key details of the significant effects; full ANOVA tables and associated statistics for all analyses can be found in our JASP notebooks located online at the following URL: https://osf.io/pefnr/.

#### Recognition Memory (Type-I)

First, we examined the influence of emotional Valence and Arousal on decision accuracy, in a two-way repeated measures ANOVA, collapsing across block and learning time conditions. We found a significant main effect of Valence (F_(1,29)_ = 9.887, η_p_2 = 0.254, p = 0.004), as well as a significant interaction between Valence and Arousal (F_(1,29)_ = 7.779, η_p_2 = 0.212, p =.009), as positive words were recognized more accurately than negative ones under low arousal conditions (T_(29)_=4.20, p_Holm_ < 0.001). Similarly, for sensitivity (d’), we found a significant effect of Valence (F_(1,29)_ = 11.34, η_p_2 = 0.28, p = 0.002), as negative words decreased d’, as well as an interaction between Valence and Arousal (F_(1,29)_= 7.34, η_p_2 = 0.20, p =0.011), as positive words were recognized more sensitively than negative ones under low arousal conditions (T_(29)_=4.304, p_Holm_ < 0.001). When analyzing the median response time, we observed a significant effect of Valence (F_(1,29)_ =0.55, η_p_2 = 0.16, p =.025), an effect of Arousal (F_(1,29)_ = 6.94, η_p_2 = 0.19, p =.013) and an interaction between these two factors (F_(1,29)_ = 7.56, η_p_2 = 0.20, p =.010), revealing that participants responded faster to positive valence under the low compared to the high arousal condition (T_(29)_=3.80, p_Holm_ =.002). Analysis of response criterion revealed no significant main effects or interactions, and no other significant effects were found (all ps >.05).

#### Metacognition (Type-II)

We then performed a second level of analysis on the metacognition data, comprising average confidence, meta-d and M-ratio. First, we performed a Valence × Arousal repeated measures ANOVA on the average confidence. This procedure showed a strong effect of Valence (F_(1,29)_ = 14.98, η_p_2 = 0.34, p < 0.001), as participants were more confident for positive valenced words. No other effects or interactions were significant. Participants were also more sensitive to their performance (Meta-d’) when responding to positive valenced words (main effect of Valence F_(1,29)_ = 11.28, η_p_2 = 0.28, p = 0.002). Concerning the M-ratio (i.e., Meta-d’/d’), which measures metacognitive efficiency, we found no main effect or interactions (all ps > 0.05). Following our pre-registered protocol, we followed up this analysis with a Bayesian ANOVA (Rouder et al., 2012) implemented in JASP (version 0.12.2) (JASP Team, 2020), to assess the strength of evidence for the null effect. This analysis compares the evidence for nested models of increasing complexity; e.g., comparing a null model to those with only main effects of valence or arousal, or a full model with main effects and interaction terms. This revealed strong relative evidence for the null overall model (including subject offsets), BFModel = 7.28; the next best model was one with a main effect of Valence whose relative BFModel = 0.74, i.e. inconclusive evidence. This analysis suggests that under the default JASP priors, it is very unlikely that Valence, Arousal, or their interaction exerted any effect on metacognitive efficiency.

#### Encoding duration

The effects of the encoding phase duration on both type I and type II task performances were not included in the pre-registration and so were assessed in an exploratory post hoc analysis. The results are described in the Supplementary Materials.

### Physiological results

#### Pulse rate variability

First, we analyzed the effect of valence and arousal on the heart rate frequency during the experimental blocks. We averaged the estimated beats per minute (Mean BPM) across the different learning times (30, 60 and 90 seconds) and submitted it to a two-way repeated measure ANOVA (Valence × Arousal). Results showed a main effect of Valence (F_(1,29)_ = 10.852, η_p_2 = 0.272, p = 0.003), meaning lower BPM for negative valenced words, but no other main or interaction effects.

For the low and high-frequency peak analysis, the peak high-frequency (HF peak) revealed an interaction between valence and arousal (F_(1,29)_ = 14.50, η_p_2 =0.33, p < 0.001) such that high-frequency cardiac oscillations were suppressed by negative emotional valence under low but not high arousal (T_(29)_=3.36, p_Holm_=.007).

Concerning the Root Mean Square of the Successive Differences (RMSSD, not shown in Fig. 4), we found a main effect of valence (F_(1,29)_ = 6.74, η_p_2 =0.19, p =.015) i.e. negative valence increased RMSSD, but no other main effect or interaction (all ps <.05).

**Fig. 4:**
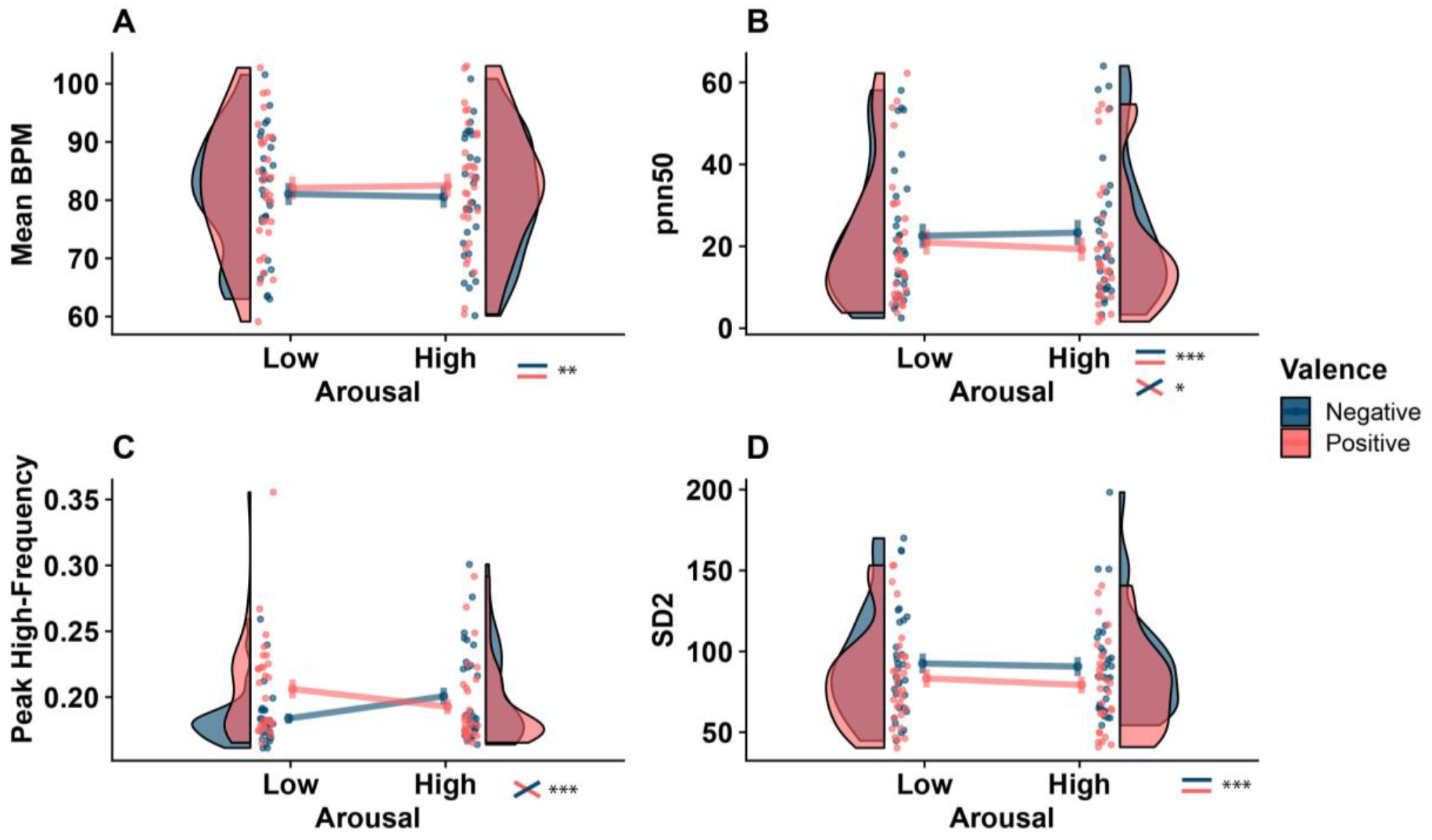
Modified raincloud plots illustrating results of pulse rate variability (PRV) analyses. PRV indices were calculated separately for each 50 trial block and averaged by condition. Mean BPM (**A**), Pnn50 (**B**), High-frequency peak (**C**), SD2 (**D**). Repeated measures ANOVA (Valence × Arousal) was then carried out for each variable separately. A significant main effect of emotional valence was observed for mean BPM, as negative valence decreased cardiac activity frequency, as well as for the pnn50 and the non-linear SD2 metric. We did not observe a main effect of Arousal, but an interaction with valence was found for the high-frequency peak, such that high-frequency cardiac oscillations were reduced by negative emotional valence under low but not high arousal. No other significant effects were found. (*** p<0.001, ** p<0.01, * p<0.05). See *Methods* and *PRVResults* for more details.

When considering the proportion of successive beat-to-beat intervals deviating by more than 50 ms (pnn50) we observed an effect of Valence (F_(1,29)_ = 24.17, η_p_2 = 0.45, p < 0.001), as well as an interaction between Valence and Arousal (F_(1,29)_ = 4.54, η_p_2 = 0.13, p =.042). Under high arousal the positive valence suppressed pnn50 while negative valence increased it (t_(29)_ =-4.98, p_Holm_ < 0.001).

Finally, we also analyzed the effect of these factors on the non-linear metrics of heart rate variability SD1 and SD2. The SD2 metric revealed an effect of Valence (F_(1,29)_ = 35.20, η_p_2 = 0.55, p < 0.001), so that negative valence increased SD2 heart rate variability, but we found no other main effects or interactions. Concerning SD1, we found no significant effects (all ps >.05). These results are illustrated in **Fig. 4**; here we reported the main significant effects, however full analyses details and results tables can be found in the HRV JASP notebook located on the Github repository for this study.

#### Event-related analysis

Next, we analyzed the time-locked instantaneous pulse rate fluctuation following word presentation. **Figure 5a** shows the evoked cardiac frequency fluctuation following the display of the two words on the screen. Following the specification of the pre-registered report, we analyzed the average of this fluctuation across time (**Figure 5b**). Here, we observed no effect of Valence, (F(1,28) = 0.441, η_p_2 = 0.015, p = 0.511), Arousal (F(1,28) = 0.003, η_p_2< 0.001, p = 0.954) or an interaction between these two factors (F(1,28) = 0.044, η_p_2 = 0.001, p =.833).

**Fig. 5:**
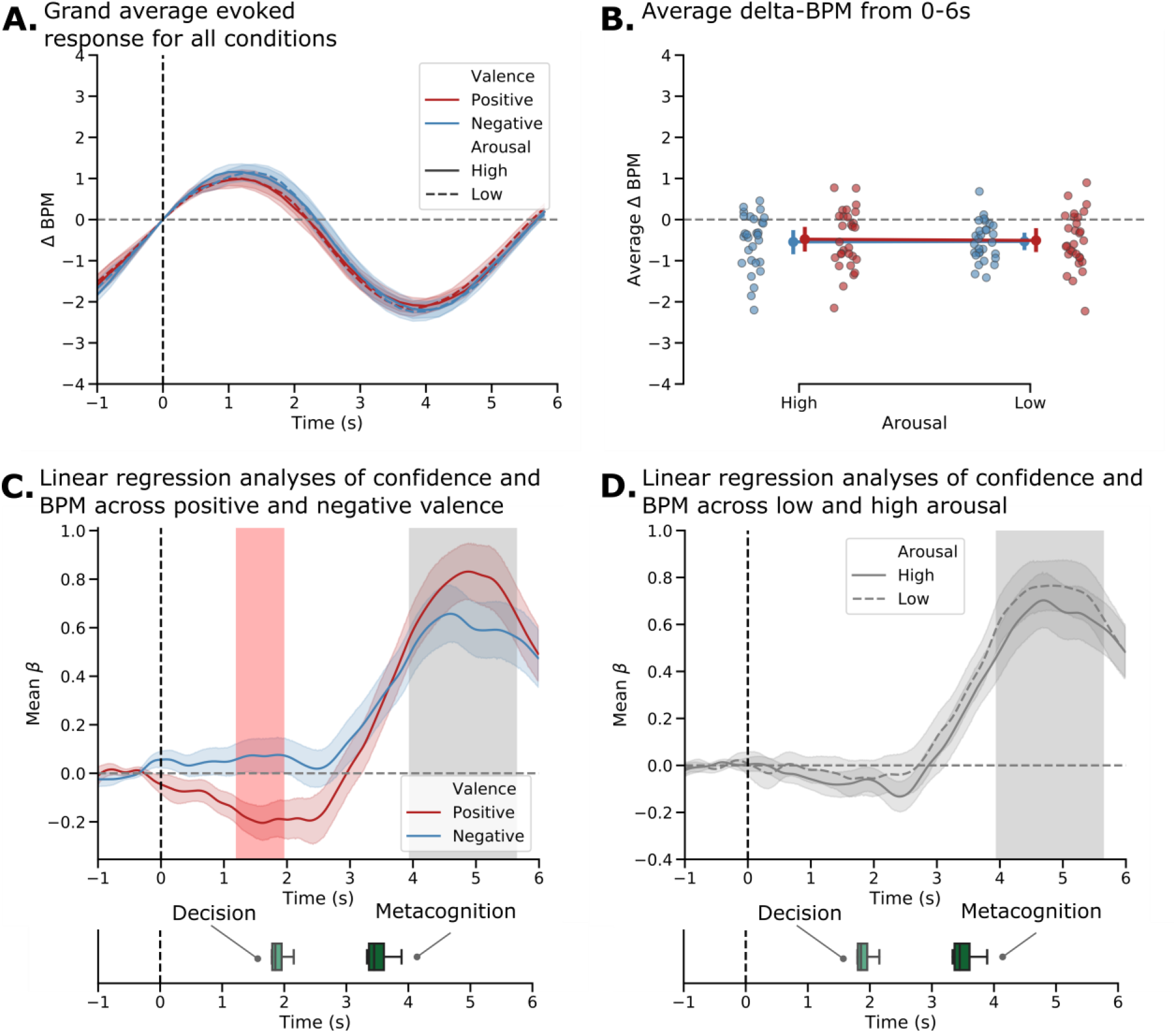
Modulation of the cardiac activity at the trial level and its relation with reported subjective confidence. **A.**Evoked pulse rate activity shows that the overall experimental procedure modulated the instantaneous cardiac frequency over time through an early acceleration component after the trial start (0-2 s) and a later deceleration component (2-6 s). This pattern is consistent with an orientation reflex, suggesting early brief integration and later sensory or memory processing. Interestingly, these two components are also time-locked with the decision and metacognition average response time. Here, we did not observe any difference between the experimental conditions. **B.**We averaged the instantaneous pulse rate in the window of interest (0-5 s) and confirmed this absence of effect and an overall diminution of cardiac frequency after the trial start. **C**. Beta values over time of the linear regression (Confidence ~ BPM) for positive and negative valence trials separately. The confidence level was associated with the instantaneous cardiac frequency during the late time window corresponding to the metacognition decision. **D.**Beta values over time of the linear regression (Confidence ~ BPM) for high and low arousal trials separately. Using the same approach, contrasting for High and Low level of arousal. Significance assessed using a cluster-level statistical permutation test (alpha=0.05). Shaded areas and error bar show the 68% CI. Significant clusters are shown by a shaded red path for condition contrast, and grey path for null tests. See online article for colour figures.

#### Linear regression

In an additional exploratory analysis, we also tested the possible interaction between the instantaneous pulse rate modulation observed during decision and metacognition and the subjective report provided by the participant. For each participant and condition separately, we used the reported confidence *C* and the instantaneous pulse rate BPM at each time point *s* of the trial *t* to fit a linear regression of the form:

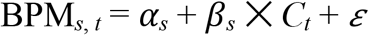

All variables were normalized and beta-values for each explanatory regressor and participant were extracted for statistical analysis. First, to test for a difference in beta values from 0 across time across all conditions, i.e. the average effect of trial by trial confidence reports on fluctuations in evoked heart-rate, replicating previous analysis linking these variables (Allen et al., 2016). We averaged between conditions assessed significance via non-parametric cluster-level t-test. Results show a significant cluster (3.94-5.65 seconds after stimulus presentation, p = 0.001). As our HRV and behavioural results emphasized an effect of stimulus valence on both metacognitive behaviour and cardiac activity, we then compared the association between confidence and instantaneous cardiac activity between different valence and arousal conditions. When comparing positive and negative valence conditions (averaging across arousal levels) we found a significant early cluster (1.20-2.96 seconds after stimulus presentation, p=0.047), suggesting that stimulus valence modulates the correlation between evoked heart-rate and confidence. Finally, we repeated this comparison for high vs. low arousal conditions, collapsing stimulus valence. This analysis found no significant clusters. See Fig. 5 for illustration of these results.

## Discussion

In this study, we investigated the influence of emotion on word memory and metamemory through a combination of experimental psychology, cognitive modelling, and physiology. We adapted a recognition memory paradigm (McCurdy et al., 2013) where participants memorised lists of words varying in arousal and valence. Stimulus valence exerted a consistent influence on recognition performance, metacognitive confidence, and physiological activity. For reaction time and accuracy, this effect was greatest for low vs. high arousal words, suggesting that the influence of stimulus valence on memory depends in part on arousal. On the physiological level, we observed an association between the subjective confidence reported by the participant and the evoked pulse rate, which was also modulated by the word valence. Our results evidence that although recognition memory is impaired for negative emotional stimuli, participants can accurately monitor and report this uncertainty. Further, monitoring the effect of emotion on memory may depend in part on integrating the associated changes in cardiac signals.

The notion that emotional events are better recalled than neutral ones has been extensively discussed (Yonelinas & Ritchey, 2015), but the psychophysiological mechanisms mediating this effect are still unclear. In some cases, emotional influences on memory have been assumed to be almost entirely ascribed to arousal, which is presumed to control the cognitive and behavioural importance of stimuli (Mather, 2007; Mather & Sutherland, 2009), while other works attributed this improvement primarily to valence (Adelman & Estes, 2013; Kensinger, 2009). Previous investigations in the perceptual domain have documented that arousing stimuli “boost” the signal-to-noise ratio of visual motion, as reflected in both models of ballistic evidence accumulation, and subjective confidence (Allen et al., 2016; Lufityanto et al., 2016). However, in our study, the effect of arousal was generally muted or dependent on stimulus valence.

Whereas on type-I task performance, stimulus valence interacted with arousal, for metacognitive type-II variables we observed only a pronounced main effect of valence with no arousal effect or interaction. In general, participant confidence reports closely matched the overall effect of stimulus emotion on performance; negative valence decreased sensitivity, increased reaction times, and decreased confidence. The robust evidence we observed for the absence of effect on metacognitive-efficiency (M-ratio) further underlines this finding; the strong null Bayes factor here demonstrates that shifts in subjective confidence were well reflected by the magnitude of any changes in type-I sensitivity, indicating that subjects make optimal use of the available memory signal during metacognitive judgements, irrespective of any conditional valence or an arousal effect. This finding suggests that, although memory is degraded under negative emotional contexts, participants can accurately account for this in their subjective confidence.

One possible explanation for the absence of arousal effect is found in our validation rating study (see Figure 2); while the valence dimension was well preserved between the ANEW database and the ratings by our participants after the task, the consistency of arousal ratings was slightly reduced. This limits the extent to which we can infer actual stimulus arousal in our data, and it may be that the stimuli were simply not sufficiently distinct or triggering for a Danish sample. Indeed, in this study arousing versus nonarousing word stimuli did not evoke a significant difference in physiological arousal response. Future studies could benefit from both a larger corpus of validated words, a more general sample of English-speaking participants, and multiple modalities of memorised stimuli which may better preserve arousal-based dimensions. Here, it should also be noted that the use of words instead of images or film is also a potential confounding factor due to the lack of vividness and complexity of the mental imagery elicited by one single item. While previous works evidenced that words classified as highly arousing and with extreme positive or negative valence are associated with better memory recall (Buchanan et al., 2006; Madan et al., 2017, 2019), other dimensions like semantic properties or the functional use of the object such as animacy are more relevant psycholinguistic dimensions to predict free recall (Madan, 2020). Again, controlling for this dimension in future studies could help to refine the influence of emotion on metamemory beyond the dimensions of valence and arousal.

In a related line of research, several investigations have linked physiological activity (e.g., as indexed by pupil dilation or cardiac acceleration) to subjective confidence and metacognition. According to influential predictive-coding accounts of metacognition (Allen et al., 2016; Meyniel et al., 2015; Moulin & Souchay, 2015), confidence reflects the width of a posterior decision variable, such that fluctuations in arousal bias the gain or precision of this distribution. Here, we examined both trial level evoked changes in cardiovascular activity and summary measures of pulse rate variability separately for each condition. When examining instantaneous heart rate variation, we observed a robust sinusoidal pattern that remained stable across conditions, similar to an orientation reflex triggered by trial onset (see Figure 5). The shape and intensity of this cardiac deceleration can reflect several cognitive processes like attention orienting, emotion processing or inhibitory control (Abercrombie et al., 2008; Critchley et al., 2005; Legrand et al., 2020). Replicating previous findings (Allen et al., 2016), we observed a robust association between trial-by-trial fluctuations in subjective confidence during this late interval, with the strength of this association being modulated by stimulus valence during the early, decision-evoked period. These results suggest that at least some variance in the monitoring of emotional inputs on metamemory could arise from monitoring the associated physiological changes. This result corroborates the notion that memory retrieval is an embodied process (Garfinkel et al., 2013), which has implications both for their conservation, the accuracy of their recall, but also their control in the case of distressing emotional memories (Gagnepain et al., 2017; Legrand et al., 2020).

Whereas no overall modulation of instantaneous heart-rate was seen for stimulus valence or arousal, here we observed substantive, robust modulations of heart rate variability (HRV) when subjects recalled negatively valenced stimuli across multiple time, frequency and non-linear indices. HRV (i.e the amount of change across time of the interbeat intervals) can reflect the influence of higher cognitive processes on cardiac frequency through the parasympathetic nervous system (Smith et al., 2017; Thayer & Lane, 2009). Across the different range of HRV indices we examined, two showed a strong valence main effect (i.e., Mean BPM & SD2), whereas others (i.e., high-frequency peak and pnn50) showed a robust interaction between these factors. Although disentangling what underlies these different effects is far from trivial, it is interesting to note the dissociation between these effects, and similarity to those observed for our type-I and type-II metamemory measures. One intriguing possibility is that the high-frequency variability indexed by the former two measures may be a more direct input for metacognitive monitoring than the others, as these showed a similar pattern of exaggerated valence effect with no effect of arousal. One means to probe this hypothesis is to correlate individual differences in the modulation of confidence by valence with each HRV metric; however, our study is underpowered for individual differences analyses (Schönbrodt & Perugini, 2013), leaving it as an intriguing avenue for future research.

Several important limitations should be considered when interpreting our HRV effects. As HRV is here calculated by collapsing across each 50 trial block, the modulations observed therein are necessarily a mixture of multiple cognitive states and perceptual inputs; future studies could benefit from disentangling the encoding, perceptual, and retrieval stages to better account for these stages of the decision process. Additionally, here we assessed heart rate variability through pulse oximetry recording. Pulse oximetry recordings are used as an alternative to the electrocardiogram (ECG) by several clinical and non-clinical studies (Quintana, Elstad, et al., 2016). The sampling rate of our device (75 Hz) is not optimal when compared to recommended standards for electrocardiogram (ECG) recording and HRV measurement (Quintana, Alvares, et al., 2016), which could limit our ability to detect true effects, particularly in the lower frequency range. Previous reports, however, have shown a strong consistency between the estimated pulse rate variability and the heart rate variability as measured through ECG (Lu et al., 2009; Schäfer & Vagedes, 2013). Similarly, we did not measure or control respiratory cycles during this study, which robustly modulate HRV measures, in particular in the lower frequency. Collectively, while our results nicely demonstrate that stimulus emotional content modulates high-frequency indices of cardiovascular arousal, future studies in this area are likely to benefit from a combination of more nuanced experimental design and a more sophisticated recording set-up.

## Conclusion

This pre-registered study sheds light on the biasing effects of valence on metamemory with possible physiological correlates of these effects. Negatively valenced stimuli globally decreased both memory performance and metacognition, supporting a role for emotions in guiding confidence and memory performance. While arousal has often been described as a possible mechanism of the beneficial effect of emotion on memory, we only found limited evidence for performance improvement under highly arousing conditions. In line with these main effects, we found that stimulus valence modulated the overall pulse rate variability and the association between instantaneous heart-rate and subjective confidence. Collectively, these results suggest that although negative stimuli do exert a degrading influence on recognition memory, participants are largely able to account for this effect in their subjective confidence, perhaps by monitoring physiological states.

## Acknowledgements

NL, CMCC, NKM, NN, and MA are supported by a Lundbeckfonden Fellowship (under Grant [R272-2017-4345]), and the AIAS-COFUND II fellowship programme that is supported by the Marie Skłodowska-Curie actions under the European Union’s Horizon 2020 (under Grant [754513]), and the Aarhus University Research Foundation.

## Data availability statement.

The project pre-registration can be found at the following link:https://osf.io/9awtb

## Data deposition and supplemental online material.

All behavioural and physiological data can be found at:https://osf.io/pefnr/

## References

Abercrombie, H., Chambers, A., Greischar, L., & Monticelli, R. (2008). Orienting, emotion, and memory: Phasic and tonic variation in heart rate predicts memory for emotional pictures in men. Neurobiol Learn Mem, 90, 644–650.

Adelman, J. S., & Estes, Z. (2013). Emotion and memory: A recognition advantage for positive and negative words independent of arousal. Cognition, 129(3), 530–535. https://doi.org/10.1016/j.cognition.2013.08.014

Allen, M., Frank, D., Schwarzkopf, D. S., Fardo, F., Winston, J. S., Hauser, T. U., & Rees, G. (2016). Unexpected arousal modulates the influence of sensory noise on confidence. ELife, 5. https://doi.org/10.7554/elife.18103

Allen, M., Poggiali, D., Whitaker, K., Marshall, T. R., & Kievit, R. A. (2019). Raincloud plots: A multi-platform tool for robust data visualization. Wellcome Open Research, 4, 63. https://doi.org/10.12688/wellcomeopenres.15191.1

Anderson, A. K. (2005). Affective Influences on the Attentional Dynamics Supporting Awareness. Journal of Experimental Psychology: General, 134(2), 258–281. https://doi.org/10.1037/0096-3445.134.2.258

Baird, B., Smallwood, J., Gorgolewski, K. J., & Margulies, D. S. (2013). Medial and Lateral Networks in Anterior Prefrontal Cortex Support Metacognitive Ability for Memory and Perception. Journal of Neuroscience, 33(42), 16657–16665. https://doi.org/10.1523/JNEUROSCI.0786-13.2013

Bradley, M. M., & Lang, P. J. (1994). Measuring emotion: The self-assessment manikin and the semantic differential. Journal of Behavior Therapy and Experimental Psychiatry, 25(1), 49–59. https://doi.org/10.1016/0005-7916(94)90063-9

Bradley, M. M., & Lang, P. J. (1999). Affective Norms for English Words (ANEW): Instruction Manual and Affective Ratings.

Buchanan, T. W., Etzel, J. A., Adolphs, R., & Tranel, D. (2006). The influence of autonomic arousal and semantic relatedness on memory for emotional words. International Journal of Psychophysiology, 61(1), 26–33. https://doi.org/10.1016/j.ijpsycho.2005.10.022

Cahill, L., & McGaugh, J. L. (1998). Mechanisms of emotional arousal and lasting declarative memory. Trends in Neurosciences, 21(7), 294–299. https://doi.org/10.1016/S0166-2236(97)01214-9

Chua, E. F., Pergolizzi, D., & Weintraub, R. R. (2014). The Cognitive Neuroscience of Metamemory Monitoring: Understanding Metamemory Processes, Subjective Levels Expressed, and Metacognitive Accuracy. In S. M. Fleming & C. D. Frith (Eds.), The Cognitive Neuroscience of Metacognition (pp. 267–291). Springer. https://doi.org/10.1007/978-3-642-45190-4_12

Critchley, H. D., Rotshtein, P., Nagai, Y., O\textquotesingleDoherty, J., Mathias, C. J., & Dolan, R. J. (2005). Activity in the human brain predicting differential heart rate responses to emotional facial expressions. Neuroimage, 24(3), 751–762. https://doi.org/10.1016/j.neuroimage.2004.10.013

Davis, R. L., & Zhong, Y. (2017). The Biology of Forgetting—A Perspective. Neuron, 95(3), 490–503. https://doi.org/10.1016/j.neuron.2017.05.039

De Martino, B., Fleming, S. M., Garrett, N., & Dolan, R. J. (2013). Confidence in value-based choice. Nature Neuroscience, 16(1), 105–110. https://doi.org/10.1038/nn.3279

Fleming, S. M. (2017). HMeta-d: Hierarchical Bayesian estimation of metacognitive efficiency from confidence ratings. Neuroscience of Consciousness, 2017(1). https://doi.org/10.1093/nc/nix007

Fleming, S. M., Dolan, R. J., & Frith, C. D. (2012). Metacognition: Computation, biology and function. Philosophical Transactions of the Royal Society B: Biological Sciences, 367(1594), 1280–1286. https://doi.org/10.1098/rstb.2012.0021

Fleming, S. M., & Lau, H. C. (2014). How to measure metacognition. Frontiers in Human Neuroscience, 8. https://doi.org/10.3389/fnhum.2014.00443

Fleming, S. M., Maniscalco, B., Ko, Y., Amendi, N., Ro, T., & Lau, H. (2015). Action-Specific Disruption of Perceptual Confidence. Psychological Science, 26(1), 89–98. https://doi.org/10.1177/0956797614557697

Fleming, S. M., Ryu, J., Golfinos, J. G., & Blackmon, K. E. (2014). Domain-specific impairment in metacognitive accuracy following anterior prefrontal lesions. Brain, 137(10), 2811–2822. https://doi.org/10.1093/brain/awu221

Gagnepain, P., Hulbert, J., & Anderson, M. C. (2017). Parallel Regulation of Memory and Emotion Supports the Suppression of Intrusive Memories. The Journal of Neuroscience, 37(27), 6423–6441. https://doi.org/10.1523/JNEUROSCI.2732-16.2017

Galvin, S. J., Podd, J. V., Drga, V., & Whitmore, J. (2003). Type 2 tasks in the theory of signal detectability: Discrimination between correct and incorrect decisions. Psychonomic Bulletin & Review, 10(4), 843–876. https://doi.org/10.3758/BF03196546

Garfinkel, S. N., Barrett, A. B., Minati, L., Dolan, R. J., Seth, A. K., & Critchley, H. D. (2013). What the heart forgets: Cardiac timing influences memory for words and is modulated by metacognition and interoceptive sensitivity: Cardiac timing and memory. Psychophysiology, 50(6), 505–512. https://doi.org/10.1111/psyp.12039

Gramfort, A. (2013). MEG and EEG data analysis with MNE-Python. Frontiers in Neuroscience, 7. https://doi.org/10.3389/fnins.2013.00267

Hauser, T. U., Allen, M., Purg, N., Moutoussis, M., Rees, G., & Dolan, R. J. (2017). Noradrenaline blockade specifically enhances metacognitive performance. ELife, 6, e24901. https://doi.org/10.7554/eLife.24901

Heyes, C., Bang, D., Shea, N., Frith, C. D., & Fleming, S. M. (2020). Knowing Ourselves Together: The Cultural Origins of Metacognition. Trends in Cognitive Sciences, 24(5), 349–362. https://doi.org/10.1016/j.tics.2020.02.007

James, W. (1884). What is an emotion? Mind, 9(34), 188–205. https://doi.org/10.1093/mind/os-ix.34.188

JASP Team. (2020). JASP (Version 0.12.2)[Computer software]. https://jasp-stats.org/

Kensinger, E. A. (2009). Remembering the Details: Effects of Emotion. Emotion Review, 1(2), 99–113. https://doi.org/10.1177/1754073908100432

Kensinger, E. A., & Corkin, S. (2004). Two routes to emotional memory: Distinct neural processes for valence and arousal. Proceedings of the National Academy of Sciences, 101(9), 3310–3315. https://doi.org/10.1073/pnas.0306408101

Kever, A., Grynberg, D., Eeckhout, C., Mermillod, M., Fantini, C., & Vermeulen, N. (2015). The body language: The spontaneous influence of congruent bodily arousal on the awareness of emotional words. Journal of Experimental Psychology: Human Perception and Performance, 41(3), 582–589. https://doi.org/10.1037/xhp0000055

Kever, A., Grynberg, D., Szmalec, A., Smalle, E., & Vermeulen, N. (2019). “Passion” versus “patience”: The effects of valence and arousal on constructive word recognition. Cognition and Emotion, 33(6), 1302–1309. https://doi.org/10.1080/02699931.2018.1561419

Kever, A., Grynberg, D., & Vermeulen, N. (2017). Congruent bodily arousal promotes the constructive recognition of emotional words. Consciousness and Cognition, 53, 81–88. https://doi.org/10.1016/j.concog.2017.06.007

Kreibig, S. D. (2010). Autonomic nervous system activity in emotion: A review. Biological Psychology, 84(3), 394–421. https://doi.org/10.1016/j.biopsycho.2010.03.010

Lau, H., & Rosenthal, D. (2011). Empirical support for higher-order theories of conscious awareness. Trends in Cognitive Sciences, 15(8), 365–373. https://doi.org/10.1016/j.tics.2011.05.009

Lee, A. L. F., Ruby, E., Giles, N., & Lau, H. (2018). Cross-Domain Association in Metacognitive Efficiency Depends on First-Order Task Types. Frontiers in Psychology, 9, 2464. https://doi.org/10.3389/fpsyg.2018.02464

Legrand, N., & Allen, M. (2020). Systole: V 0.1.2—September 2020 (0.1.2) [Computer software]. Zenodo. https://doi.org/10.5281/ZENODO.3607912

Legrand, N., Etard, O., Vandevelde, A., Pierre, M., Viader, F., Clochon, P., Doidy, F., Peschanski, D., Eustache, F., & Gagnepain, P. (2020). Long-term modulation of cardiac activity induced by inhibitory control over emotional memories. Scientific Reports, 10(1), 15008. https://doi.org/10.1038/s41598-020-71858-2

Lipponen, J. A., & Tarvainen, M. P. (2019). A robust algorithm for heart rate variability time series artefact correction using novel beat classification. Journal of Medical Engineering & Technology, 43(3), 173–181. https://doi.org/10.1080/03091902.2019.1640306

Lu, G., Yang, F., Taylor, J. A., & Stein, J. F. (2009). A comparison of photoplethysmography and ECG recording to analyse heart rate variability in healthy subjects. Journal of Medical Engineering & Technology, 33(8), 634–641. https://doi.org/10.3109/03091900903150998

Lufityanto, G., Donkin, C., & Pearson, J. (2016). Measuring Intuition: Nonconscious Emotional Information Boosts Decision Accuracy and Confidence. Psychological Science, 27(5), 622–634. https://doi.org/10.1177/0956797616629403

Macmillan, N. A., & Creelman, C. D. (2004). Detection Theory: A User’s Guide (2nd ed.). Psychology Press. https://doi.org/10.4324/9781410611147

Madan, C. R. (2020). Exploring word memorability: How well do different word properties explain item free-recall probability? Psychonomic Bulletin & Review. https://doi.org/10.3758/s13423-020-01820-w

Madan, C. R., Scott, S. M. E., & Kensinger, E. A. (2019). Positive emotion enhances association-memory. Emotion, 19(4), 733–740. https://doi.org/10.1037/emo0000465

Madan, C. R., Shafer, A. T., Chan, M., & Singhal, A. (2017). Shock and awe: Distinct effects of taboo words on lexical decision and free recall. Quarterly Journal of Experimental Psychology (2006), 70(4), 793–810. https://doi.org/10.1080/17470218.2016.1167925

Maniscalco, B., & Lau, H. (2012). A signal detection theoretic approach for estimating metacognitive sensitivity from confidence ratings. Consciousness and Cognition, 21(1), 422–430. https://doi.org/10.1016/j.concog.2011.09.021

Mather, M. (2007). Emotional Arousal and Memory Binding: An Object-Based Framework. Perspectives on Psychological Science, 2(1), 33–52. https://doi.org/10.1111/j.1745-6916.2007.00028.x

Mather, M., & Sutherland, M. (2009). Disentangling the Effects of Arousal and Valence on Memory for Intrinsic Details. Emotion Review, 1(2), 118–119. https://doi.org/10.1177/1754073908100435

Mazancieux, A., Fleming, S. M., Souchay, C., & Moulin, C. J. A. (2020). Is there a G factor for metacognition? Correlations in retrospective metacognitive sensitivity across tasks. Journal of Experimental Psychology: General. https://doi.org/10.1037/xge0000746

McCurdy, L. Y., Maniscalco, B., Metcalfe, J., Liu, K. Y., de Lange, F. P., & Lau, H. (2013). Anatomical Coupling between Distinct Metacognitive Systems for Memory and Visual Perception. Journal of Neuroscience, 33(5), 1897–1906. https://doi.org/10.1523/JNEUROSCI.1890-12.2013

Meyniel, F., Sigman, M., & Mainen, Z. F. (2015). Confidence as Bayesian Probability: From Neural Origins to Behavior. Neuron, 88(1), 78–92. https://doi.org/10.1016/j.neuron.2015.09.039

Montefinese, M., Ambrosini, E., Fairfield, B., & Mammarella, N. (2014). The adaptation of the Affective Norms for English Words (ANEW) for Italian. Behavior Research Methods, 46(3), 887–903. https://doi.org/10.3758/s13428-013-0405-3

Moors, A., De Houwer, J., Hermans, D., Wanmaker, S., van Schie, K., Van Harmelen, A.-L., De Schryver, M., De Winne, J., & Brysbaert, M. (2013). Norms of valence, arousal, dominance, and age of acquisition for 4, 300 Dutch words. Behavior Research Methods, 45(1), 169–177. https://doi.org/10.3758/s13428-012-0243-8

Morales, J., Lau, H., & Fleming, S. M. (2018). Domain-General and Domain-Specific Patterns of Activity Supporting Metacognition in Human Prefrontal Cortex. The Journal of Neuroscience, 38(14), 3534–3546. https://doi.org/10.1523/JNEUROSCI.2360-17.2018

Moulin, C., & Souchay, C. (2015). An active inference and epistemic value view of metacognition. Cognitive Neuroscience, 6(4), 221–222. https://doi.org/10.1080/17588928.2015.1051015

Ochsner, K. N. (2000). Are affective events richly recollected or simply familiar? The experience and process of recognizing feelings past. Journal of Experimental Psychology: General, 129(2), 242–261. https://doi.org/10.1037/0096-3445.129.2.242

Otgaar, H., Howe, M. L., Patihis, L., Merckelbach, H., Lynn, S. J., Lilienfeld, S. O., & Loftus, E. F. (2019). The Return of the Repressed: The Persistent and Problematic Claims of Long-Forgotten Trauma. Perspectives on Psychological Science, 14(6), 1072–1095. https://doi.org/10.1177/1745691619862306

Peirce, J., Gray, J. R., Simpson, S., MacAskill, M., Höchenberger, R., Sogo, H., Kastman, E., & Lindeløv, J. K. (2019). PsychoPy2: Experiments in behavior made easy. Behavior Research Methods, 51(1), 195–203. https://doi.org/10.3758/s13428-018-01193-y

Quintana, D. S., Alvares, G. A., & Heathers, J. A. J. (2016). Guidelines for Reporting Articles on Psychiatry and Heart rate variability (GRAPH): Recommendations to advance research communication. Translational Psychiatry, 6(5), e803–e803. https://doi.org/10.1038/tp.2016.73

Quintana, D. S., Elstad, M., Kaufmann, T., Brandt, C. L., Haatveit, B., Haram, M., Nerhus, M., Westlye, L. T., & Andreassen, O. A. (2016). Resting-state high-frequency heart rate variability is related to respiratory frequency in individuals with severe mental illness but not healthy controls. Scientific Reports, 6(1), 37212. https://doi.org/10.1038/srep37212

Redondo, J., Fraga, I., Padrón, I., & Comesaña, M. (2007). The Spanish adaptation of ANEW (Affective Norms for English Words). Behavior Research Methods, 39(3), 600–605. https://doi.org/10.3758/BF03193031

Reggev, N., Zuckerman, M., & Maril, A. (2011). Are all judgments created equal?: An fMRI study of semantic and episodic metamemory predictions. Neuropsychologia, 49(5), 1332–1342. https://doi.org/10.1016/j.neuropsychologia.2011.01.013

Rollwage, M., Loosen, A., Hauser, T. U., Moran, R., Dolan, R. J., & Fleming, S. M. (2020). Confidence drives a neural confirmation bias. Nature Communications, 11(1), 2634. https://doi.org/10.1038/s41467-020-16278-6

Rouder, J. N., Morey, R. D., Speckman, P. L., & Province, J. M. (2012). Default Bayes factors for ANOVA designs. Journal of Mathematical Psychology, 56(5), 356–374. https://doi.org/10.1016/jjmp.2012.08.001

Schäfer, A., & Vagedes, J. (2013). How accurate is pulse rate variability as an estimate of heart rate variability? International Journal of Cardiology, 166(1), 15–29. https://doi.org/10.1016/j.ijcard.2012.03.119

Schönbrodt, F. D., & Perugini, M. (2013). At what sample size do correlations stabilize? Journal of Research in Personality, 47(5), 609–612. https://doi.org/10.1016/j.jrp.2013.05.009

Shaffer, F., & Ginsberg, J. P. (2017). An Overview of Heart Rate Variability Metrics and Norms. Frontiers in Public Health, 5, 258. https://doi.org/10.3389/fpubh.2017.00258

Shaffer, F., McCraty, R., & Zerr, C. L. (2014). A healthy heart is not a metronome: An integrative review of the heart’s anatomy and heart rate variability. Frontiers in Psychology, 5. https://doi.org/10.3389/fpsyg.2014.01040

Shea, N., Boldt, A., Bang, D., Yeung, N., Heyes, C., & Frith, C. D. (2014). Supra-personal cognitive control and metacognition. Trends in Cognitive Sciences, 18(4), 186–193. https://doi.org/10.1016/j.tics.2014.01.006

Shields, G. S., Sazma, M. A., McCullough, A. M., & Yonelinas, A. P. (2017). The effects of acute stress on episodic memory: A meta-analysis and integrative review. Psychological Bulletin, 143(6), 636–675. https://doi.org/10.1037/bul0000100

Smith, R., Thayer, J. F., Khalsa, S. S., & Lane, R. D. (2017). The hierarchical basis of neurovisceral integration. Neuroscience & Biobehavioral Reviews, 75, 274–296. https://doi.org/10.1016/j.neubiorev.2017.02.003

Tauber, S. K., & Dunlosky, J. (2012). Can older adults accurately judge their learning of emotional information? Psychology and Aging, 27(4), 924–933. https://doi.org/10.1037/a0028447

Thayer, J. F., & Lane, R. D. (2009). Claude Bernard and the heart-brain connection: Further elaboration of a model of neurovisceral integration. Neuroscience and Biobehavioral Reviews, 33(2), 81–88. https://doi.org/10.1016/j.neubiorev.2008.08.004

Valenza, G., Citi, L., Lanatá, A., Scilingo, E. P., & Barbieri, R. (2014). Revealing Real-Time Emotional Responses: A Personalized Assessment based on Heartbeat Dynamics. Scientific Reports, 4(1). https://doi.org/10.1038/srep04998

Vallat, R. (2018). Pingouin: Statistics in Python. Journal of Open Source Software, 3(31), 1026. https://doi.org/10.21105/joss.01026

van Gent, P., Farah, H., van Nes, N., & van Arem, B. (2019). Analysing Noisy Driver Physiology Real-Time Using Off-the-Shelf Sensors: Heart Rate Analysis Software from the Taking the Fast Lane Project. Journal of Open Research Software, 7, 32. https://doi.org/10.5334/jors.241

Ye, Q., Zou, F., Lau, H., Hu, Y., & Kwok, S. C. (2018). Causal Evidence for Mnemonic Metacognition in Human Precuneus. The Journal of Neuroscience, 38(28), 6379–6387. https://doi.org/10.1523/JNEUROSCI.0660-18.2018

Yeung, N., & Summerfield, C. (2012). Metacognition in human decision-making: Confidence and error monitoring. Philosophical Transactions of the Royal Society B: Biological Sciences, 367(1594), 1310–1321. https://doi.org/10.1098/rstb.2011.0416

Yonelinas, A. P., & Ritchey, M. (2015). The slow forgetting of emotional episodic memories: An emotional binding account. Trends in Cognitive Sciences, 19(5), 259–267. https://doi.org/10.1016/j.tics.2015.02.009

