## Supplementary material for "Emotional Metacognition: Stimulus Valence Modulates Cardiac Arousal and Metamemory"

In this section, we analyse the behavioural data reported in the manuscript by including the Learning time parameter. For each level of the Arousal*Valence interaction, each participant underwent 3 blocks of 50 words with different learning time (30, 60 or 90 seconds). This period controlled the duration of the initial presentation of the words, which influenced the strength of encoding. Here, we report the values of the variables of interest reported in Figure 3 for each learning time condition separately.

The code used to extract the data can be found in the 5 – LearningTime(Supplementary Material).ipynb file. The repeated measure ANOVA were conducted using JASP (v. 0.14), and can be found in the learningTime_analysis.jasp file. In the following figures, error bar represent the 68% CI.

### 1. Response time

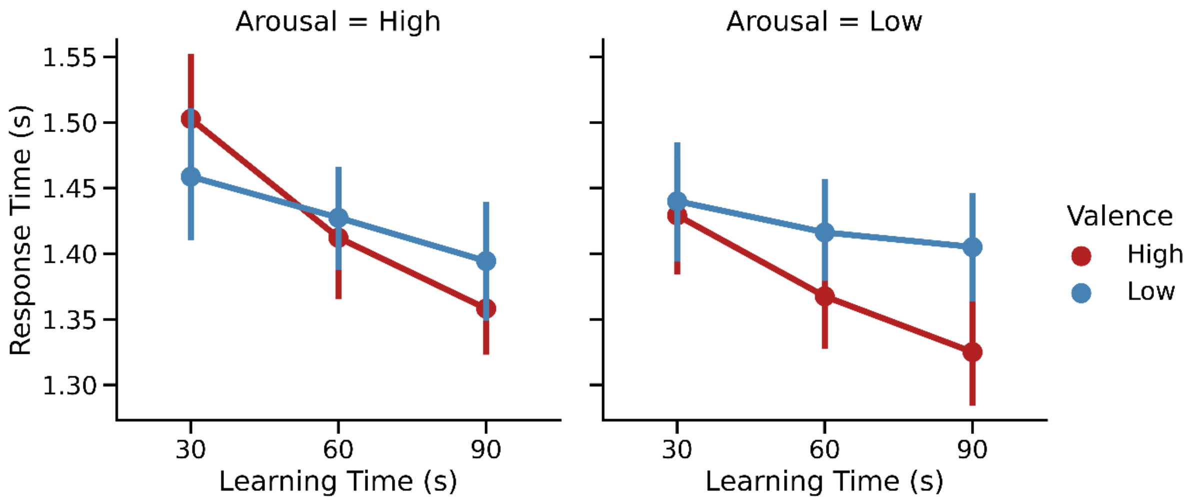


We examined the influence of emotional Valence, Arousal and learning time on response time using a three-factor repeated measures ANOVA. We found a significant effect of **Arousal** ($F_{(1,34)}$ = 7.557, $\eta_p 2$= 0.182, p = 0.009), **Learning time** ($F_{(2,68)}$ = 6.969, $\eta_p 2$= 0.170, p = 0.002), as well as a significant interaction between **Valence** and **Learning Time** ($F_{(2,68)}$ = 3.835, $\eta_p 2$= 0.101, p = .026).

**Within Subjects Effects**

| Cases | Sum of Squares | df | Mean Square | F | p | $\eta^2_p$ |
|---|---|---|---|---|---|---|
| Valence | 0.143 | 1 | 0.143 | 1.764 | 0.193 | 0.049 |
| Residuals | 2.759 | 34 | 0.081 | | | |
| Arousal | 0.077 | 1 | 0.077 | 1.727 | 0.198 | 0.048 |
| Residuals | 1.512 | 34 | 0.044 | | | |
| Learning Time | 0.886 | 2 | 0.443 | 3.644 | 0.031 | 0.097 |
| Residuals | 8.266 | 68 | 0.122 | | | |
| Valence ✻ Arousal | 0.025 | 1 | 0.025 | 0.434 | 0.515 | 0.013 |
| Residuals | 1.922 | 34 | 0.057 | | | |
| Valence ✻ Learning Time | 0.312 | 2 | 0.156 | 4.289 | 0.018 | 0.112 |
| Residuals | 2.475 | 68 | 0.036 | | | |
| Arousal ✻ Learning Time | 0.302 | 2 | 0.151 | 3.452 | 0.037 | 0.092 |
| Residuals | 2.971 | 68 | 0.044 | | | |
| Valence ✻ Arousal ✻ Learning Time | 0.197 | 2 | 0.099 | 1.619 | 0.206 | 0.045 |
| Residuals | 4.142 | 68 | 0.061 | | | |

*Note.* Type III Sum of Squares

[a] Mauchly's test of sphericity indicates that the assumption of sphericity is violated (p < .05).

#### 2. Criterion

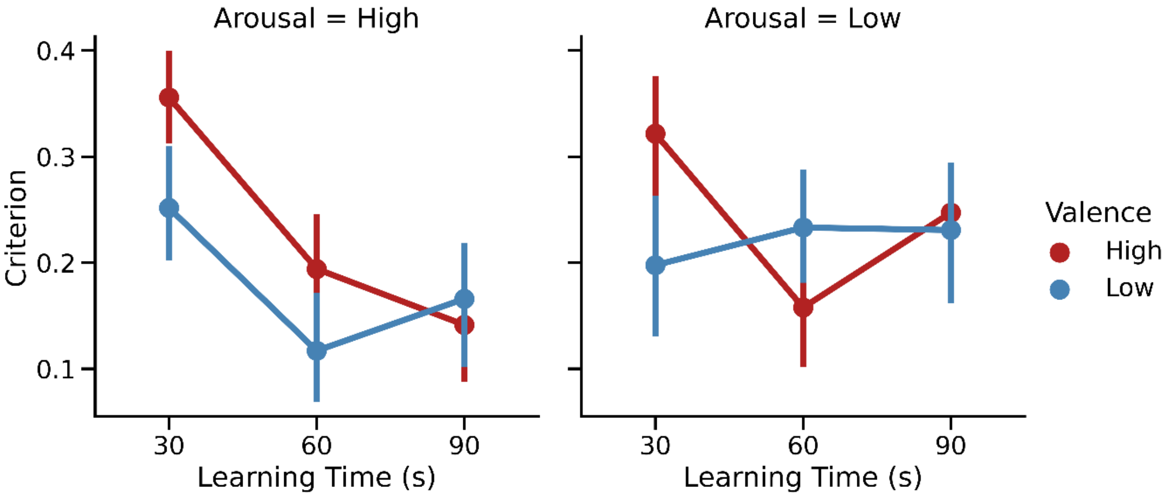


We examined the influence of emotional Valence, Arousal and learning time on criterion using a three-factor repeated measures ANOVA. We found a significant effect of **Learning time** ($F_{(2,68)}$ = 3.644, $\eta_p$2= 0.097, p = 0.031), as well as an interaction between **Learning time** and **Valence** ($F_{(2,68)}$ = 4.289, $\eta_p$2= 0.112, p = 0.018), and between **Learning time** and **Arousal** ($F_{(2,68)}$ = 3.452, $\eta_p$2= 0.037, p = 0.092).

**Within Subjects Effects**

| Cases | Sum of Squares | df | Mean Square | F | p | $\eta^2_p$ |
|---|---|---|---|---|---|---|
| Valence | 0.037 | 1 | 0.037 | 7.348 | 0.010 | 0.178 |
| Residuals | 0.171 | 34 | 0.005 | | | |
| Arousal | 7.258e -5 | 1 | 7.258e -5 | 0.018 | 0.894 | 5.313e -4 |
| Residuals | 0.137 | 34 | 0.004 | | | |
| Learning Time | 1.188 | 2 | 0.594 | 81.624 | < .001 | 0.706 |
| Residuals | 0.495 | 68 | 0.007 | | | |
| Valence ✻ Arousal | 0.030 | 1 | 0.030 | 6.908 | 0.013 | 0.169 |
| Residuals | 0.146 | 34 | 0.004 | | | |
| Valence ✻ Learning Time | 0.043 | 2 | 0.022 | 4.820 | 0.011 | 0.124 |
| Residuals | 0.307 | 68 | 0.005 | | | |
| Arousal ✻ Learning Time | 0.016 | 2 | 0.008 | 1.917 | 0.155 | 0.053 |

| | | | | | | |
|---|---|---|---|---|---|---|
| Residuals | 0.281 | 68 | 0.004 | | | |
| Valence ∗ Arousal ∗ Learning Time | 0.012 | 2 | 0.006 | 1.408 | 0.252 | 0.040 |
| Residuals | 0.287 | 68 | 0.004 | | | |

*Note.* Type III Sum of Squares
[a] Mauchly's test of sphericity indicates that the assumption of sphericity is violated (p < .05).

#### 3. Accuracy

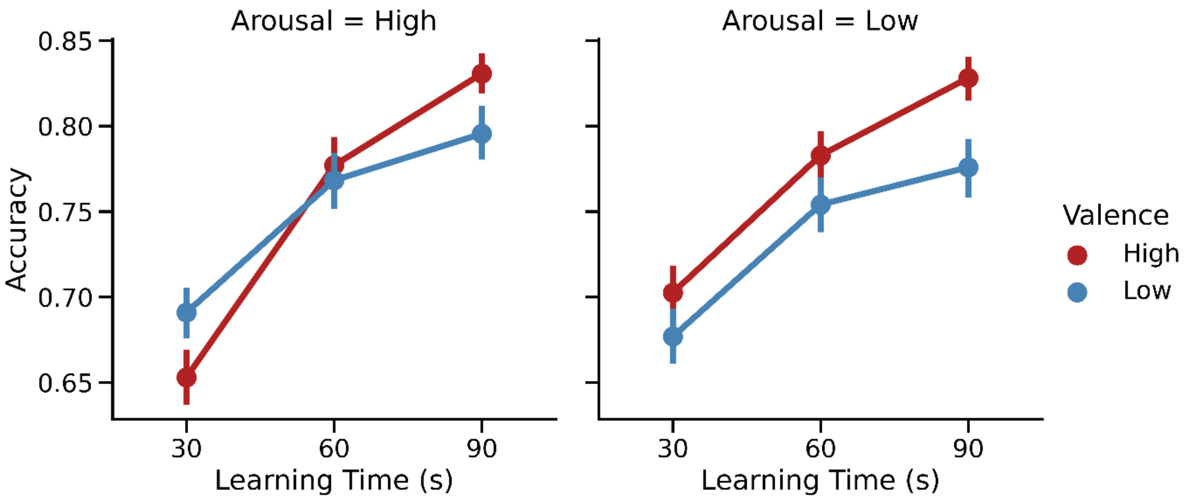


We examined the influence of emotional Valence, Arousal and learning time on response accuracy using a three-factor repeated measures ANOVA. We found a significant effect of **Valence** ($F_{(1,34)}$ = 7.348, $\eta_p 2$= 0.178, p = 0.010), of **Learning Time** ($F_{(2,68)}$ = 81.624, $\eta_p 2$= 0.706, p < 0.001), as well as an interaction between **Arousal** and **Valence** ($F_{(1,34)}$ = 6.908, $\eta_p 2$= 0.169, p = 0.013), and between **Learning time** and **Valence** ($F_{(2,68)}$ = 4.820, $\eta_p 2$= 0.124, p = 0.011).

**Within Subjects Effects**

| Cases | Sum of Squares | df | Mean Square | F | p | $\eta^2_p$ |
|---|---|---|---|---|---|---|
| Valence | 6.607 | 1 | 6.607 | 16.443 | < .001 | 0.326 |
| Residuals | 13.663 | 34 | 0.402 | | | |
| Arousal | 0.266 | 1 | 0.266 | 1.244 | 0.273 | 0.035 |
| Residuals | 7.261 | 34 | 0.214 | | | |
| Learning Time | 116.075 | 2 | 58.037 | 76.187 | < .001 | 0.691 |

| | | | | | | |
|---|---|---|---|---|---|---|
| Residuals | 51.801 | 68 | 0.762 | | | |
| Valence ∗ Arousal | 0.893 | 1 | 0.893 | 5.085 | 0.031 | 0.130 |
| Residuals | 5.971 | 34 | 0.176 | | | |
| Valence ∗ Learning Time | 2.518 | 2 | 1.259 | 4.862 | 0.011 | 0.125 |
| Residuals | 17.607 | 68 | 0.259 | | | |
| Arousal ∗ Learning Time | 0.260 | 2 | 0.130 | 0.574 | 0.566 | 0.017 |
| Residuals | 15.403 | 68 | 0.227 | | | |
| Valence ∗ Arousal ∗ Learning Time | 0.223 | 2 | 0.111 | 0.541 | 0.585 | 0.016 |
| Residuals | 13.999 | 68 | 0.206 | | | |

*Note.* Type III Sum of Squares
[a] Mauchly's test of sphericity indicates that the assumption of sphericity is violated (p < .05).

#### 4. Confidence

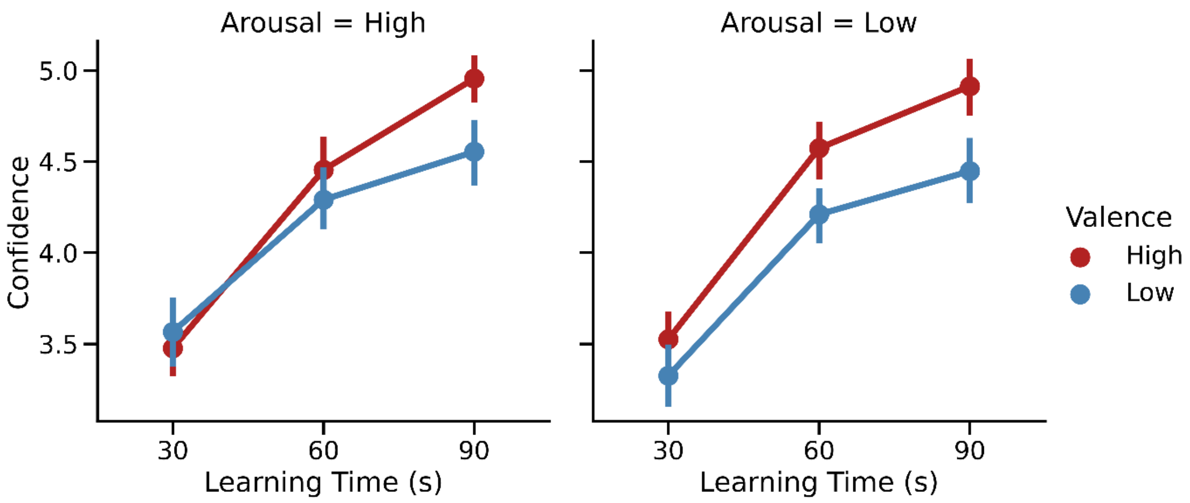


We examined the influence of emotional Valence, Arousal and learning time on response confidence using a three-factor repeated measures ANOVA. We found a significant effect of **Valence** ($F_{(1,34)}$ = 16.443, $\eta_p2$= 0.326, p < 0.001), of **Learning Time** ($F_{(2,68)}$ = 76.187, $\eta_p2$= 0.691, p < 0.001), as well as an interaction between **Arousal** and **Valence** ($F_{(1,34)}$ = 5.085, $\eta_p2$= 0.130, p = 0.031), and between **Valence** and **Learning time** ($F_{(2,68)}$ = 4.862, $\eta_p2$= 0.125, p = 0.011).

**Within Subjects Effects**

| Cases | Sum of Squares | df | Mean Square | F | p | $\eta^2_p$ |
|---|---|---|---|---|---|---|
| Valence | 6.607 | 1 | 6.607 | 16.443 | < .001 | 0.326 |
| Residuals | 13.663 | 34 | 0.402 | | | |
| Arousal | 0.266 | 1 | 0.266 | 1.244 | 0.273 | 0.035 |
| Residuals | 7.261 | 34 | 0.214 | | | |
| Learning Time | 116.075 | 2 | 58.037 | 76.187 | < .001 | 0.691 |
| Residuals | 51.801 | 68 | 0.762 | | | |
| Valence ✻ Arousal | 0.893 | 1 | 0.893 | 5.085 | 0.031 | 0.130 |
| Residuals | 5.971 | 34 | 0.176 | | | |
| Valence ✻ Learning Time | 2.518 | 2 | 1.259 | 4.862 | 0.011 | 0.125 |
| Residuals | 17.607 | 68 | 0.259 | | | |
| Arousal ✻ Learning Time | 0.260 | 2 | 0.130 | 0.574 | 0.566 | 0.017 |
| Residuals | 15.403 | 68 | 0.227 | | | |
| Valence ✻ Arousal ✻ Learning Time | 0.223 | 2 | 0.111 | 0.541 | 0.585 | 0.016 |
| Residuals | 13.999 | 68 | 0.206 | | | |

*Note.* Type III Sum of Squares

[a] Mauchly's test of sphericity indicates that the assumption of sphericity is violated (p < .05).

## 5. d'

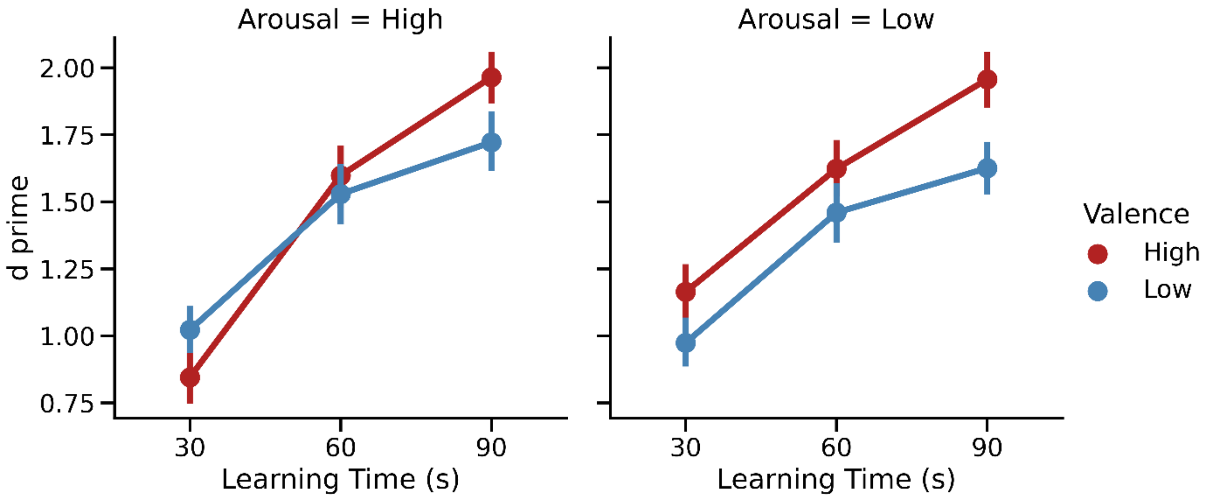


We examined the influence of emotional Valence, Arousal and learning time on *d'* using a three-factor repeated measures ANOVA. We found a significant effect of **Valence** ($F_{(1,34)}$ = 9.343, $\eta_p 2$= 0.216, p = 0.004), of **Learning Time** ($F_{(2,68)}$ = 74.5577, $\eta_p 2$= 0.687, p < 0.001), as well as an interaction between **Arousal** and **Valence** ($F_{(1,34)}$ = 4.394, $\eta_p 2$= 0.114, p = 0.044).

**Within Subjects Effects**

| Cases | Sum of Squares | df | Mean Square | F | p | $\eta^2_p$ |
|---|---|---|---|---|---|---|
| Valence | 1.958 | 1 | 1.958 | 9.343 | 0.004 | 0.216 |
| Residuals | 7.124 | 34 | 0.210 | | | |
| Arousal | 0.043 | 1 | 0.043 | 0.254 | 0.618 | 0.007 |
| Residuals | 5.755 | 34 | 0.169 | | | |
| Learning Time | 48.552 | 2 | 24.276 | 74.557 | < .001 | 0.687 |
| Residuals | 22.141 | 68 | 0.326 | | | |
| Valence ✻ Arousal | 0.885 | 1 | 0.885 | 4.394 | 0.044 | 0.114 |
| Residuals | 6.850 | 34 | 0.201 | | | |
| Valence ✻ Learning Time | 1.394 | 2 | 0.697 | 2.969 | 0.058 | 0.080 |
| Residuals | 15.967 | 68 | 0.235 | | | |

| | | | | | | |
|---|---|---|---|---|---|---|
| Arousal ∗ Learning Time | 0.708 | 2 | 0.354 | 1.840 | 0.167 | 0.051 |
| Residuals | 13.074 | 68 | 0.192 | | | |
| Valence ∗ Arousal ∗ Learning Time | 0.444 | 2 | 0.222 | 1.148 | 0.323 | 0.033 |
| Residuals | 13.155 | 68 | 0.193 | | | |

*Note.* Type III Sum of Squares
[a] Mauchly's test of sphericity indicates that the assumption of sphericity is violated (p < .05).

#### 6. Meta-d'

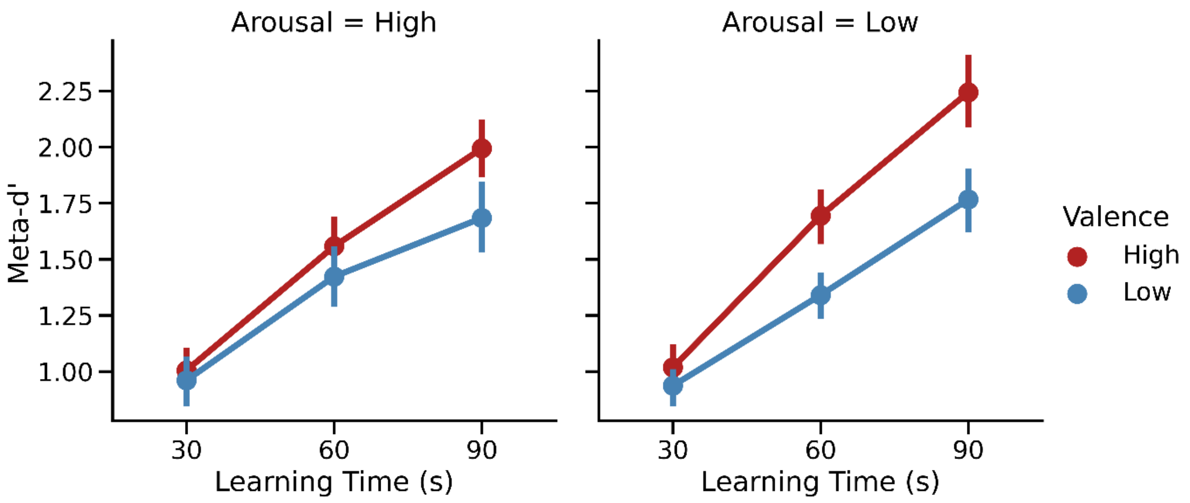


We examined the influence of emotional Valence, Arousal and learning time on meta-*d'* using a three-factor repeated measures ANOVA. We found a significant effect of **Valence** ($F_{(1,34)} = 15.979$, $\eta_p 2 = 0.320$, $p < 0.001$), of **Learning Time** ($F_{(2,68)} = 94.418$, $\eta_p 2 = 0.735$, $p < 0.001$), as well as an interaction between **Learning Time** and **Valence** ($F_{(1,34)} = 3.501$, $\eta_p 2 = 0.093$, $p = 0.036$).

**Within Subjects Effects**

| Cases | Sum of Squares | df | Mean Square | F | p | $\eta^2_p$ |
|---|---|---|---|---|---|---|
| Valence | 5.728 | 1 | 5.728 | 15.979 | < .001 | 0.320 |
| Residuals | 12.187 | 34 | 0.358 | | | |
| Arousal | 0.411 | 1 | 0.411 | 1.946 | 0.172 | 0.054 |

| | | | | | | |
|---|---|---|---|---|---|---|
| Residuals | 7.179 | 34 | 0.211 | | | |
| Learning Time | 62.354 | 2 | 31.177 | 94.418 | < .001 | 0.735 |
| Residuals | 22.454 | 68 | 0.330 | | | |
| Valence ∗ Arousal | 0.520 | 1 | 0.520 | 2.407 | 0.130 | 0.066 |
| Residuals | 7.352 | 34 | 0.216 | | | |
| Valence ∗ Learning Time | 1.922 | 2 | 0.961 | 3.501 | 0.036 | 0.093 |
| Residuals | 18.668 | 68 | 0.275 | | | |
| Arousal ∗ Learning Time | 0.585 | 2 | 0.292 | 1.022 | 0.365 | 0.029 |
| Residuals | 19.446 | 68 | 0.286 | | | |
| Valence ∗ Arousal ∗ Learning Time | 0.153 | 2 | 0.076 | 0.415 | 0.662 | 0.012 |
| Residuals | 12.518 | 68 | 0.184 | | | |

*Note*. Type III Sum of Squares

[a] Mauchly's test of sphericity indicates that the assumption of sphericity is violated (p < .05).

### 7. M-ratio

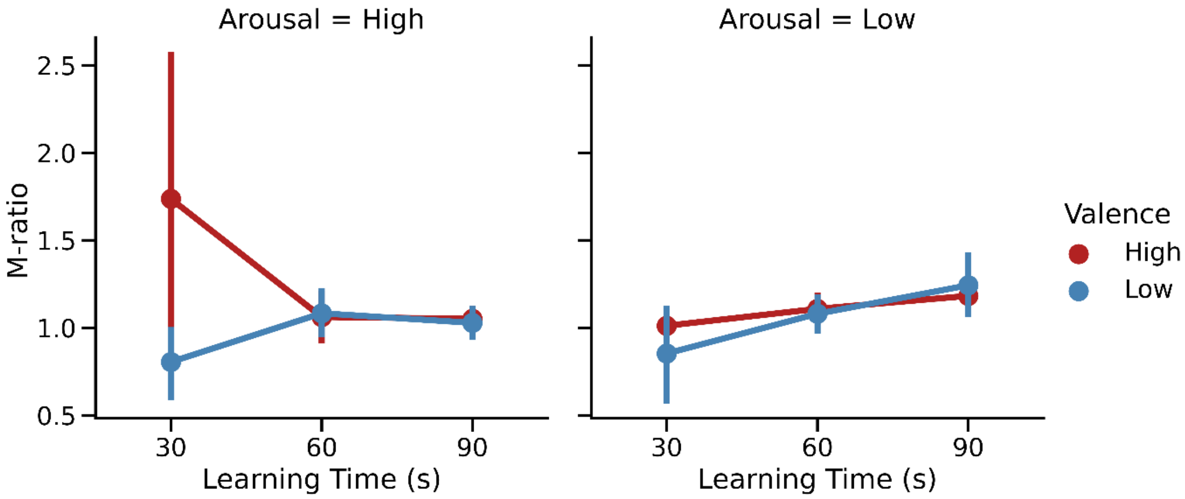


We examined the influence of emotional Valence, Arousal and learning time on m-ratio using a three-factor repeated measures ANOVA. Here, we found no significant effect of any of these factors. It should be noted that this analysis results in very few trials per bin, which can strongly bias the non-hierarchical Mratio fits, resulting in large negative or positive values.

**Within Subjects Effects**

| Cases | Sum of Squares | df | Mean Square | F | p | $\eta^2_p$ |
|---|---|---|---|---|---|---|
| Valence | 3.283 | 1 | 3.283 | 2.070 | 0.159 | 0.057 |
| Residuals | 53.917 | 34 | 1.586 | | | |
| Arousal | 0.241 | 1 | 0.241 | 0.135 | 0.715 | 0.004 |
| Residuals | 60.477 | 34 | 1.779 | | | |
| Learning Time | 0.130[a] | 2[a] | 0.065[a] | 0.027[a] | 0.974[a] | 7.861e-4 |
| Residuals | 165.448 | 68 | 2.433 | | | |
| Valence ∗ Arousal | 1.905 | 1 | 1.905 | 0.721 | 0.402 | 0.021 |
| Residuals | 89.852 | 34 | 2.643 | | | |
| Valence ∗ Learning Time | 7.150[a] | 2[a] | 3.575[a] | 1.236[a] | 0.297[a] | 0.035 |
| Residuals | 196.726 | 68 | 2.893 | | | |

| | | | | | | |
|---|---|---|---|---|---|---|
| Arousal ✻ Learning Time | 4.827 [a] | 2 [a] | 2.414 [a] | 0.715 [a] | 0.493 [a] | 0.021 |
| Residuals | 229.510 | 68 | 3.375 | | | |
| Valence ✻ Arousal ✻ Learning Time | 3.413 [a] | 2 [a] | 1.706 [a] | 0.898 [a] | 0.412 [a] | 0.026 |
| Residuals | 129.224 | 68 | 1.900 | | | |

*Note.* Type III Sum of Squares

[a] Mauchly's test of sphericity indicates that the assumption of sphericity is violated ($p < .05$).
